# Genome-wide Comparative Analysis Reveals Possible Common Ancestors of NBS Domain Containing Genes in Hybrid *Citrus sinensis* Genome and Original *Citrus clementina* Genome

**DOI:** 10.1101/008219

**Authors:** Yunsheng Wang, Lijuan Zhou, Dazhi Li, Amy Lawton-Rauh, Pradip K. Srimani, Liangying Dai, Yongping Duan, Feng Luo

**Author notes:** Corresponding authors: Liangying Dai: College of Plant Protection, Hunan Agricultural University, Changsha, 410128, China Feng Luo: 310 McAdams Hall, Clemson University, Clemson, SC 29634-0974, USA.

## Abstract

**Background:** Recently available whole genome sequences of three citrus species: one *Citrus clementina* and two *Citrus sinensis* genomes have made it possible to understand the features of candidate disease resistance genes with nucleotide-binding sites (NBS) domain in Citrus and how NBS genes differ between hybrid and original Citrus species.

**Result:** We identified and re-annotated NBS genes from three citrus genomes and found similar numbers of NBS genes in those citrus genomes. Phylogenetic analysis of all citrus NBS genes across three genomes showed that there are three approximately evenly numbered groups: one group contains the Toll-Interleukin receptor (TIR) domain and two different groups that contain the Coiled Coil (CC) domain. Motif analysis confirmed that the two groups of CC-containing NBS genes are from different evolutionary origins. We partitioned NBS genes into clades using NBS domain sequence distances and found most clades include NBS genes from all three citrus genomes. This suggests that NBS genes in three citrus genomes may come from shared ancestral origins. We also mapped the re-sequenced reads of three pomelo and three Mandarin orange genomes onto the *Citrus sinensis* genome. We found that most NBS genes of the hybrid *C. sinensis* genome have corresponding homologous genes in both pomelo and mandarin genome. The homologous NBS genes in pomelo and mandarin may explain why the NBS genes in their hybrid *Citrus sinensis* are similar to those in *Citrus clementina* in this study. Furthermore, sequence variation amongst citrus NBS genes were shaped by multiple independent and shared accelerated mutation accumulation events among different groups of NBS genes and in different citrus genomes.

**Conclusion:** Our comparative analyses yield valuable insight into the understanding of the structure, evolution and organization of NBS genes in *Citrus* genomes. There are significantly more NBS genes in *Citrus* genomes compared to other plant species. NBS genes in hybrid *C. sinensis* genomes are very similar to those in progenitor *C. clementina* genome and they may be derived from possible common ancestral gene copies. Furthermore, our comprehensive analysis showed that there are three groups of plant NBS genes while CC-containing NBS genes can be divided into two groups.

## Background

Disease resistance genes (R genes) are essential components of plant immune systems. Amongst five diverse classes of disease resistance genes [1], the largest class is those proteins that have nucleotide-binding site (NBS) domain and Leucine-Rich Repeat (LRR) domain. The NBS domain is evolutionarily conserved and is typically used to identify and characterize plant R genes. The LRR domains in R genes that mediate direct or indirect interactions [2, 3] with pathogen molecules are usually rapidly evolving to adapt to the change of pathogen ligands [4]. NBS domain-containing R genes were classified into two major types based on their domain structures in N terminus: proteins with a Toll-Interleukin receptor (TIR) domain and proteins with a Coiled Coil (CC) domain. While the TIR domain is usually well defined [5], the CC domain has higher variation and is less well characterized.

Genome wide analyses of NBS genes in many genomes have showed that NBS R genes are diverse in number, structure and organization [5–12]. For example, grass genomes studied to date do not have TIR NBS genes; on the other hand, dicot genomes usually contain more TIR NBS genes than CC NBS genes [13]. Meanwhile, comparative analyses of NBS genes have facilitated the understanding of the organization, classification and evolution of NBS R genes. Comparison of R genes in multiple genomes indicated that R genes are shaped by dynamic birth-and-death processes [14]. Comparative analyses of NBS genes in two Arabidopsis genomes (*A. thaliana* and *A. lyrata*) indicate that mating system shift from outcrossing to inbreeding has had a limited impact on the numbers of NBS genes, at least in short term[15]. Comparative analyses of NBS genes of diploid *Phaseolus* and tetraploid *Glycine* species concluded that whole genome duplication did not result in NBS R gene number increases. Recently, comparison of R genes in diverse grass genomes by Yang *et al.* showed that rapid evolution in R genes in *Zea*, *Sorghum*, and *Brachypodium* is associated with rice blast disease resistance [16].

Citrus species are amongst the most important fruit trees and have been cultivated for more than 4000 years [17, 18]. Phylogenetic analyses using molecular markers showed that other cultivated citrus species, such as, sweet orange, grapefruit, and lemon, are derived from three original cultivated *Citrus* species: *Citrus medica* (citron), *Citrus reticulata* (Mandarin orange) and *Citrus maxima* (pomelo) [19, 20]. For example, the *Citrus sinensis* (sweet orange) is suggested to be the backcross hybrid of *C. maxima* (pomelo) and *Citrus reticulata* (Mandarin orange) [19–21]. Phylogenetic analyses of partial NBS genes of *Poncirus trifoliata* (trifoliate orange), *Citrus reticulata* (tangerine) and their F1 progeny showed that NBS genes of *Poncirus trifoliata* (trifoliate orange), *Citrus reticulata* formed genus-specific clades. Additionally, NBS genes of their F1 progeny had sister relationships to only one of the parents [22]. This suggests that NBS genes in crossing hybrid citrus species are also different from those in original citrus species.

Recently available draft whole genome sequences of three citrus species: *Citrus clementina*, *Citrus sinensis* from USA and *Citrus sinensis* from China [21], have made it possible to scan and identify all NBS genes in those genomes. In this study, we performed a genome-wide comparative analysis of NBS genes in three citrus genomes (*Citrus clementina*, *Citrus sinensis* from China and *Citrus sinensis* from USA) to address the following questions: (1) What are the features of NBS genes in Citrus, such as numbers, physical locations, within-gene domain structures, and evolutionary dynamics of NBS genes?; (2) Do NBS genes differ between hybrid and non-hybrid *Citrus* species?; and (3) Do *Citrus* NBS genes differ from NBS genes in other plant genomes?

## Results

### Identification and classification of *Citrus* NBS Genes

We searched the *Citrus clementina*, *Citrus sinensis* USA and *Citrus sinensis* China genomes for genes containing the NBS domain using hmmsearch. Then, the NBS-containing genes were confirmed through homology searches against the Swiss protein database (see “Materials and Methods”). We identified similar numbers of NBS domain-containing genes amongst these three genomes. We found 618, 650 and 508 NBS genes from *Citrus clementina*, *Citrus sinensis* China and *Citrus sinensis* USA genomes respectively (Table 1). Among them, 413, 499 and 484 NBS genes were predicted in the original gene annotations and 205, 151 and 24 NBS genes in *Citrus clementina*, *Citrus sinensis* China and *Citrus sinensis* USA respectively were newly predicted in this project (Table S1).

**Table 1.** Classification of citrus NBS genes

| Class | *Citrus clementina* | *Citrus sinensis* China | *Citrus sinensis* USA |
| --- | --- | --- | --- |
| CNL | 117 | 113 | 60 |
| TNL | 82 | 77 | 30 |
| NL | 68 | 62 | 85 |
| Others | 351 | 398 | 333 |
| Total | 618 | 650 | 508 |

The CNL class of NBS genes contains three domains: Coiled Coil (CC), nucleotide-binding sites (NBS) and Leucine-Rich Repeat (LRR). The TNL class of NBS genes contains three domains: Toll-Interleukin receptor (TIR), nucleotide-binding sites (NBS) and Leucine-Rich Repeat (LRR). The NL class of NBS genes contains two domains: nucleotide-binding sites (NBS) and Leucine-Rich Repeat (LRR). The other NBS genes contain no LRR domain.

NBS genes could be classified into different classes based on their domain structures [23]. We searched the Toll-Interleukin receptor (TIR) and Coiled Coil (CC) domains in the N-terminal region and the Leucine-Rich Repeat (LRR) domains in the C-terminal of NBS genes. We identified 117 NBS genes with CC-NBS-LRR (CNL) domains, 82 NBS genes with TIR-NBS-LRR (TNL) domains, 68 NBS genes with NBS-LRR (NL) and 351 others NBS genes without LRR domains (N, CN and TN) from *Citrus clementina* (Table 1). We also identified 113 CNL, 77 TNL, 68 NL and 398 others NBS genes from *Citrus sinensis* China and 60 CNL, 30 TNL, 85 NL and 333 others NBS genes from *Citrus sinensis* USA (Table 1). In comparison to other genomes, there are many more *Citrus* NBS genes without the LRR domain. There are only 43.2%, 38.8% and 34.4% NBS genes with existing LRR domains in *Citrus clementina*, *Citrus sinensis* China and *Citrus sinensis* USA, respectively, while 72% NBS genes in Arabidopsis [23] and 78% NBS genes in *Populus trichocarpa* [24] have LRR domain.

The structures of TNL and CNL NBS genes are significantly different. The TNL NBS genes tend to have more introns than that of CNL NBS genes as previously found in *Arabidopsis* [23] and *Populus* [24]. The average numbers of introns in TNL NBS genes are 4.39, 4.70 and 4.47, while the numbers of introns in CNL NBS genes are 0.89, 1.15 and 1.38 in *Citrus clementina*, *Citrus sinensis* China and *Citrus sinensis* USA respectively (Figure S1). The median numbers of introns of different types of NBS genes in the three *Citrus* genomes are similar: four in TNL NBS genes and one in CNL NBS genes.

The NBS genes were unevenly distributed in the *Citrus* scaffolds/chromosomes. There were 217, 158 and 110 NBS genes, totaling 78.3% of 618 NGS genes, distributed in scaffolds 5, 3, and 7 of *C. clementina,* respectively. There were 125, 121 and 75 NBS genes distributed in chromosomes 1, 3 and 5 of *C. sinensis* China. The majority (95 out of 107) of NBS genes with TIR domain in *C. clementina* was distributed in scaffold 3. There were 33 and 51 (out of 102) NBS genes with TIR domain in *C. sinensis* China distributed on chromosome 5 and chromosome unknown, respectively.

### Phylogenetic and Clade Analysis of the *Citrus* NBS Genes

We used the protein sequences of NBS domains, which are the most conserved part of NBS genes, to construct a phylogenetic tree of NBS genes. We only selected NBS domain sequences longer than 200 amino acid residues and contain both P-loop and MHDV motifs. Finally, 442 *Citrus clementina*, 393 *Citrus sinensis* China, and 264 *Citrus sinensis* USA NBS domain sequences were used for phylogenetic analysis (Table S1). A Maximum Likelihood (ML) phylogenetic tree of these 1,099 NBS genes was then constructed using FastTree [25]. As shown in Figure 1, the un-rooted phylogenetic tree can be divided into three main groups. Most NBS genes containing the TIR domain are in one branch and most NBS genes containing a CC domain comprise the other two branches (Figure 1). Therefore, we denoted the three branches as TIR, CC1 and CC2.

There were 452, 382 and 265 *Citrus* NBS genes in CC1, CC2 and TIR groups, respectively. Table S2 lists the group classification of 1,099 *Citrus* NBS genes. For each sub-group, we constructed a new ML phylogenetic tree rooted by *Streptomyces coelicolor* protein P25941 (Figure S2). The NBS domains in CC groups, especially in the CC1 group, were relatively more diverged in sequence versus those in TIR group. The average Poisson corrected distances between sequence pairs within each group were 0.947, 0.827 and 0.663 for CC1, CC2 and TIR groups, respectively. We aligned the NBS domains sequences in each group using mafft with default parameters (--auto). The average percentages of identities for NBS domain sequences were 40%, 44% and 52% for CC1, CC2 and TIR groups, respectively.

**Figure 1.**
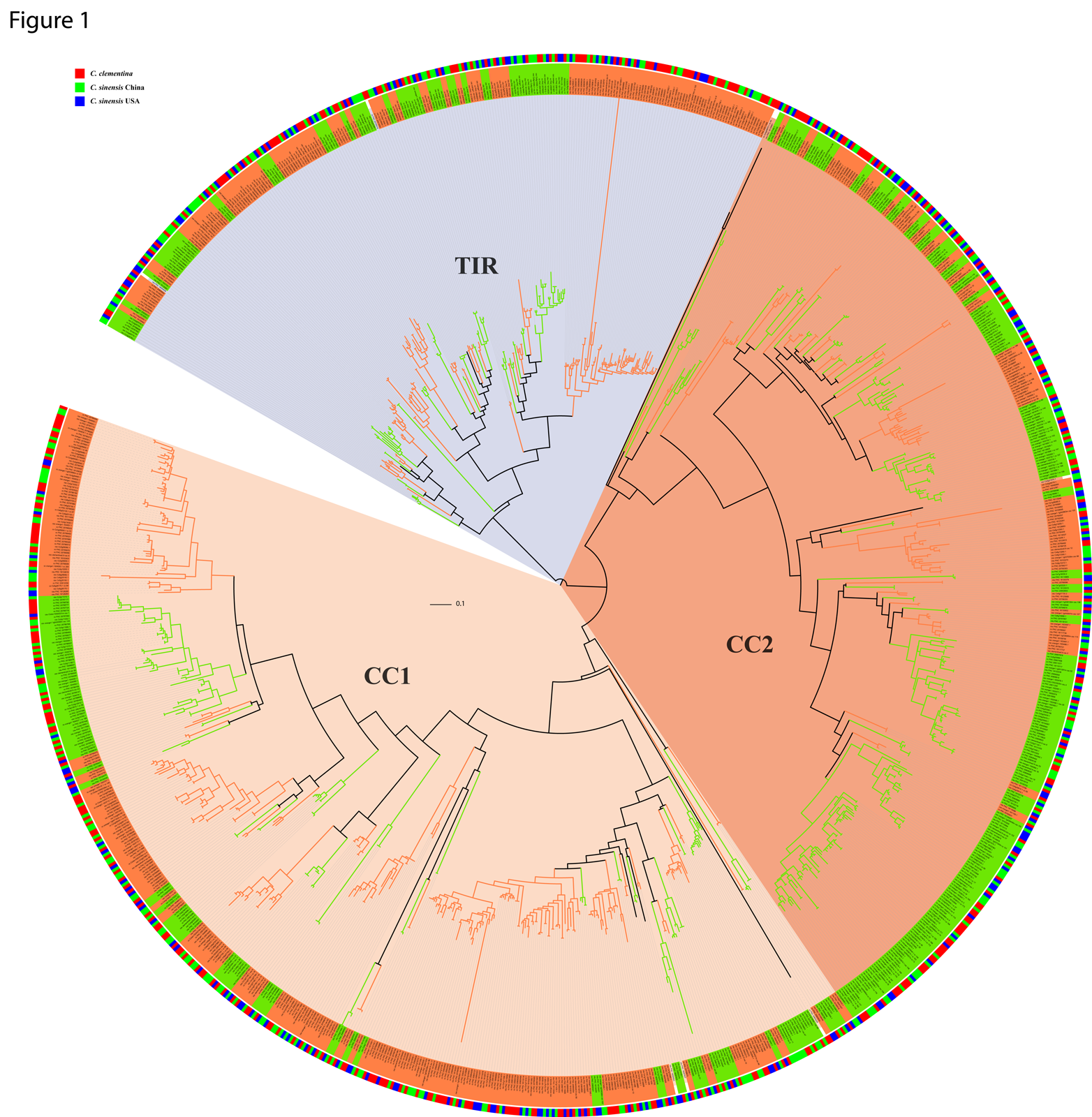
**Maximum likelihood phylogenetic tree of *Citrus* NBS-LRR genes constructed from multiple sequences alignment of NBS domain**. There are three groups in the phylogenetic tree: CC1, CC2 and TIR. Clades were classified using PhyloPart as shown in alternating color. The outer circle shows species with *Citrus clementina* (Cc) in red, *Citrus sinensis* China (CsCN) in green and *Citrus sinensis* USA (CsUSA) in blue.

**Figure 2.**
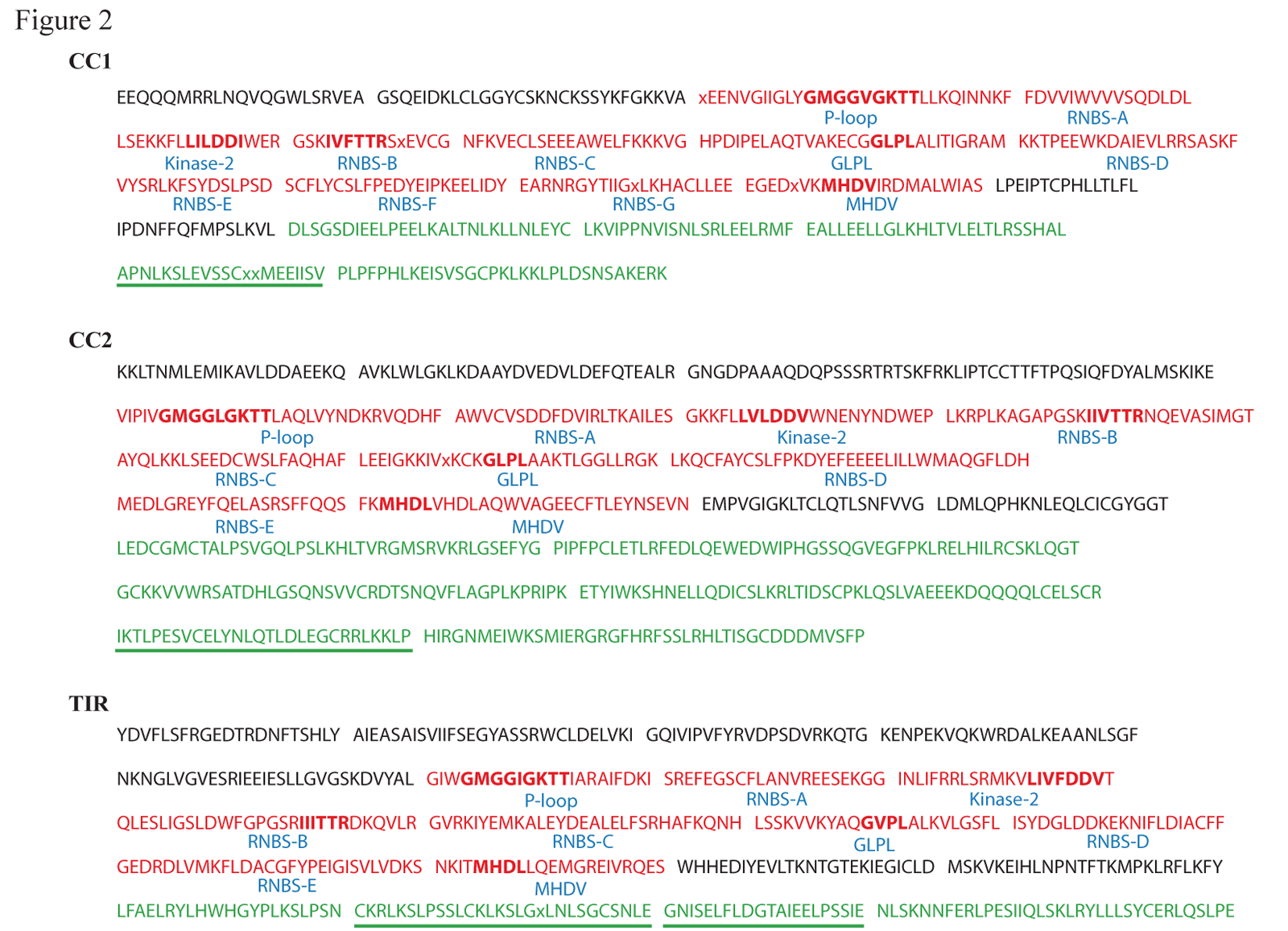
**Architecture of NBS gene motifs in *Citrus*.** Top 20 motifs identified by MEME for each group of citrus NBS genes are listed. The motifs underlined in green were repeated in C-terminal (LRR domain).

The number of LRR domains varied amongst the three classes of *Citrus* NBS genes. The number of LRR domains in the TIR group is significantly higher than the numbers of LRR domains in the CC1 and CC2 groups (Figure S3). The majority of CC1 NBS genes have only one LRR domain but the majority of TIR NBS genes have 2 LRR domains. The most frequent type of LRR domain in the CC1 group is LRR_8 (PF13855), but LRR_1 (PF00560) is the most frequent domain in both the CC2 group and the TIR group (Table S3).

Previous studies showed that NBS genes from different species fall into separate phylogenetic groups [26]. In contrast, NBS genes in the three *Citrus* genomes we investigated are mixed amongst branches of phylogenetic tree as shown in the Figure 1. We partitioned the NBS gene tree into clades based on NBS domain sequence distance using PhyloPart with distance threshold of 0.025. This resulted in 114 clades (with more than two genes in each clade) and eight orphan genes (Table S2). The numbers of clades in each NBS gene group are similar. There are 39, 42, 32 clades in CC1, CC2 and TIR groups, respectively. We calculated average sequence similarities between NBS domains in each clade using alistat in SQUID (http://selab.janelia.org/software.html). The minimum average sequence similarities of NBS domains in each clade was 70%. The largest clade included 90 NBS genes. On average, there were 9.56 genes per clade. There were seven clades with more than 40 genes. Among these clades, four clades were in CC1; two clades were in CC2 and one clade was in the TIR group. Most of the clades contain NBS genes from the three *Citrus* genomes (Figure 1, Table 2). There are 89 clades with more than three NBS genes and 87 of these clades contain members from each of the three *Citrus* genomes. Together, these results imply that all of the NBS genes of the three *Citrus* genomes may have possible common ancestors.

**Table 2.** Type of citrus NBS gene clades

| NBS genes from citrus genomes | Number of Clade |
| --- | --- |
| *Citrus clementina*, *Citrus sinensis* China, *Citrus sinensis* USA | 87 |
| *Citrus clementina*, *Citrus sinensis* China | 13 |
| *Citrus clementina*, *Citrus sinensis* USA | 7 |
| *Citrus sinensis* China, *Citrus sinensis* USA | 5 |
| *Citrus clementina*, | 1 |
| *Citrus sinensis* China, | 0 |
| *Citrus sinensis* USA | 1 |

To understand the evolutionary dynamics maintaining functional constraint while maintaining so many gene family members, we tested for evidence of accelerated sequence evolution (positive selection) by searching for positively-selected sites within each clade. We detected positively selected sites in 53.5% (61) of clades. There were 18, 24 and 19 clades with sites under positive selection in CC1, CC2 and TIR groups, respectively. In total, we detected 541 positively selected sites. Consistent with previous reports [5], there are more positively selected sites in the C-terminal region of NBS genes (LRR domains) than those in the N-terminal and NBS domains (Table S4). Approximately 56.4% (305 out of 541) of the positively selected sites were located in the C-terminal region (LRR domain). While most clades have positively selected sites in the LRR domain, there were clades with a greater number of positively selected sites within the NBS domain. For example, there were 46 positively selected sites in Clade_1260 that are located in NBS domain while there are only two positively selected sites in LRR domain (Figure S4).

### Mapping Re-sequencing Reads Showed the Conservation of NBS Genes in *Citrus* Genomes

To verify that *Citrus sinensis* (sweet orange) is a backcross hybrid of *C. maxima* (pomelo) and *C. reticulata* (Mandarin orange) [21], Xu et al. re-sequenced three *C. maxima* cultivars and three *C. reticulata* cultivars and showed that SSR and SNP markers in the *C. sinensis* genome derive from the *C. maxima* and *C. reticulata* genomes at an approximately ratio of 1:3 [21]. Since our comparison of *C. maxima* and *C. reticulata* genomes showed that most of their NBS genes are very similar in sequence, we investigated the relationships amongst the NBS genes of *C. sinensis* (sweet orange), *C. maxima* (pomelo), and *C. reticulata* (Mandarin orange) to see if the evolutionary dynamics of NBS genes is informative.

We mapped the re-sequenced reads of three *Citrus maxima* (pomelo) genomes and three *C. reticulata* (Mandarin orange) genomes to hybridized genome of *Citrus sinensis* (sweet orange) China using BWA with mem method [27] and calculated the coverage of each NBS gene in *Citrus sinensis* China. We detected 524 of 650 *Citrus sinensis* China NBS genes with more than 50% coverage in all of the six resequenced genomes. Additionally, 249 of 650 *Citrus sinensis* China NBS genes with more than 85% coverage in all six resequenced genomes and 592 (91%) of *Citrus sinensis* China NBS genes with coverage over 85% in at least one of the resequenced *Citrus* genomes. Using a cutoff of 15% coverage as missing calls over the entire length of the gene, only 25 (<4%) of *Citrus sinensis* China NBS genes seem to be deleted in at least one of the six resequenced genomes. No NBS genes were lost amongst all of the six resequenced genomes.

Resequenced reads tended to map to exons rather than to introns (Table S5). For example, the entire coding region of NBS gene Cs9g13310 was 3,693 bp in length and covered by all reads from the resequenced *Citrus* samples. However, the 16 kb introns were not consistently present in all of the resequenced reads. When we used exon presence instead of whole gene coverage, 403 out of 650 (62%) *Citrus sinensis* China NBS genes have more than 85% coverage in all six resequenced *Citrus* genomes and 645 (99%) *Citrus sinensis* China NBS genes have more than 85% coverage in at least one of the resequenced *Citrus* genomes. Most *Citrus sinensis* China NBS genes have high exon coverage on both the *C. maxima* (pomelo) and *C. reticulata* (Mandarin orange) genomes (Figure S5). There were only three *Citrus sinensis* China NBS genes, Cs1g02140.1, Cs6g02120.1 and Cs7g02220.1, with low (<40%) exon coverage in all three *C. maxima* genomes, but high coverage was achieved in at least two of the three *C. reticulata* genomes (Figure S5). Our mapping results showed that most NBS genes of *C. sinensis* (sweet orange) have a corresponding copy in the resequenced *C. maxima* (pomelo) and *C. reticulata* (Mandarin orange) genomes.

### Motif Patterns in *Citrus* NBS Genes

To further examine the gene structures of three *Citrus* NBS gene groups, we searched the motifs in the sequences of each group using MEME. Figure 2 lists the top 20 motifs identified by MEME from the CC1, CC2 and TIR *Citrus* NBS gene groups.

The motifs in the N-terminal of CC1 and CC2 groups showed little similarity to each other. The MEME found two motifs from CC1 group. One of them, EEQQQMRRLNQVQGWLSRVEA, was present in 356 of 452 CC1 NBS genes. The other motif, GSQEIDKLCLGGYCSKNCKSSYKFGKKVA, was present in 189 of 452 CC1 NBS genes. Both motifs have high level of sequence similarities with the CC NBS genes of *Arabidopsis* (Figure S6(A)). The MEME found three motifs from CC2 group. Two of the motifs, KKLTNMLEMIKAVLDDAEEKQ and AVKLWLGKLKDAAYDVEDVLDEFQTEALR were identified in 325 and 340 of 382 CC2 NBS genes, respectively. The motif AVKLWLGKLKDAAYDVEDVLDEFQTEALR has high levels of similarity with the motif identified from the CC NBS genes of *Oryza sativa japonica* (Japonica rice) [28] (Figure S6 (B)). The different motifs in CC1 and CC2 NBS genes implied that they may be from different evolutionary origins.

For the TIR group, the MEME identified five motifs in the N-terminal. They can be found in 76% to 90% of 265 TIR *Citrus* NBS genes. The first four motifs YDVFLSFRGEDTRDNFTSHLY, AIEASAISVIIFSEGYASSRWCLDELVKI, GQIVIPVFYRVDPSDVRKQTG, and ENPEKVQKWRDALKEA were very similar to the TIR1-4 motifs identified from TIR NBS genes in *Arabidopsis* [23] and *Populus trichocarpa* [24] (Figure S6 (C)).

The MEME algorithm identified eight motifs from NBS domains of CC2 and TIR groups, which are similar to the motif structures of NBS domains in *Arabidopsis* [23] and *Populus trichocarpa* NBS genes [24]. MEME results also showed that the motif structure of NBS domain of CC1 groups is slightly different from those of CC2 and TIR groups. Ten motifs were identified from NBS domains of CC1 group with two extra RNBS motifs (RNBS-F, RNBS-G) between GLPL and MHDV motifs. Five of eight motifs: P-loop, Kinase-2, RNBS-B, GLPL and MHDV, showed high levels of sequence similarity amongst the three NBS groups, which were also similar to those motifs from *Arabidopsis* [23] and *Populus trichocarpa* NBS genes [24] . The MHDV motif in *Citrus* was often slightly modified to MHDL, as found in *Arabidopsis* and *Populus* previously. Meanwhile, the motifs RNBS-A, RNBS-C, and RNBS-D were quite dissimilar to each other amongst three groups.

The LRR domains in the C-terminal of NBS genes usually have high sequence diversity as they play a role in recognizing pathogen virulence proteins. The MEME algorithm identified five, six, and four motifs from LRR domains of CC1, CC2 and TIR NBS genes. Most of the motifs contained LxxL repeats. Motifs from different groups were highly variable in sequence, which implied that the three groups of NBS genes have different levels of sequence, and thus functional, constraint and play different roles in the *Citrus* immune system. Some motifs had repeated several times in the same gene. For examples, motif APNLKSLEVSSCxxMEEIISV was found in 367 of 452 CC1 group with an average 4.8 motifs per gene; motif IKTLPESVCELYNLQTLDLEGCRRLKKLP was found in 332 of 382 CC2 group with an average 2.8 times per gene and motifs CKRLKSLPSSLCKLKSLGxLNLSGCSNLE and GNISELFLDGTAIEELPSSIE were found in 208 and 209 of 265 TIR group with 2.5 and 2.1 times per gene, respectively.

### Analysis of NBS Gene Clusters in *Citrus* Genomes

The majority of *Citrus* NBS genes were physically clustered in genome (Table 3). 525 of 618 NBS genes in *C. clementina* were found in 108 clusters and 500 of 650 NBS genes in *C. sinensis* China were found in 126 clusters (Table S2). Although the assembly of *C. sinensis* USA is more fragmentary, there still are 207 of 508 NBS genes present in 72 clusters. The largest number of gene clusters in C*. clementina, C. sinensis* China and *C. sinensis* USA contain 55, 37 and 13 NBS genes, respectively.

Most clusters contain NBS genes from the same group. In *C. clementina*, there were 38 clusters with NBS genes of CC1 group, 40 clusters with NBS genes of CC3 group and 23 clusters with NBS genes of TIR group. Only seven out of 108 clusters contain the NBS genes from two or three groups. In *C. sinensis* China, there were 41 clusters with NBS genes of CC1 group, 44 clusters with NBS genes of CC2 group and 26 clusters with NBS genes of TIR group. There were 15 clusters containing NBS genes from two or three groups in *C. sinensis* China. In *C. sinensis* USA, there were 27 clusters with NBS genes of CC1 group, 25 clusters with NBS genes of CC2 group and 15 clusters with NBS genes of TIR group.

**Table 3.** Citrus NBS genes in clusters

| Citrus species | Total NBS genes | Number of NBS genes in cluster | Number of clusters | Percent of NBS genes in cluster (%) |
| --- | --- | --- | --- | --- |
| *Citrus clementina* | 618 | 525 | 108 | 84.9 |
| *Citrus sinensis* China | 650 | 500 | 126 | 76.9 |
| *Citrus sinensis* USA | 508 | 207 | 72 | 40.7 |

Only five clusters contain NBS genes from different groups in *C. sinensis* USA. The lower number of clusters in *C. sinensis* USA may due to its fragmental assembly.

The sequences of NBS genes within clusters are much more similar to each other than those between clusters (T-test p value < 2.2e-16). The mean identities between genes within and between clusters were 0.628 and 0.393 respectively (Figure 3). Furthermore, NBS genes in the same cluster tend to be in the same strand, which indicates that the NBS genes in the clusters are due to tandem duplication. We detected 204 pairs of tandem duplications in *C. clementina* and 217 pairs of tandem duplications in *C. sinensis* China using MCScanX [29]. The numbers of tandem duplications within CC1 and CC2 groups are much greater than that within the TIR group. Among 204 pairs in *C. clementina*, 90 and 85 pairs were present in the CC1 and CC2 groups and only 29 pairs from TIR group. Among 214 tandem gene pairs in *C. sinensis* China, 88 and 87 pairs were present in the CC1 and CC2 groups and only 42 pairs in the TIR group. The fewer tandem duplications of NBS genes in the TIR group may be the reason that there are fewer clusters of TIR groups in *Citrus* genomes.

**Figure 3.**
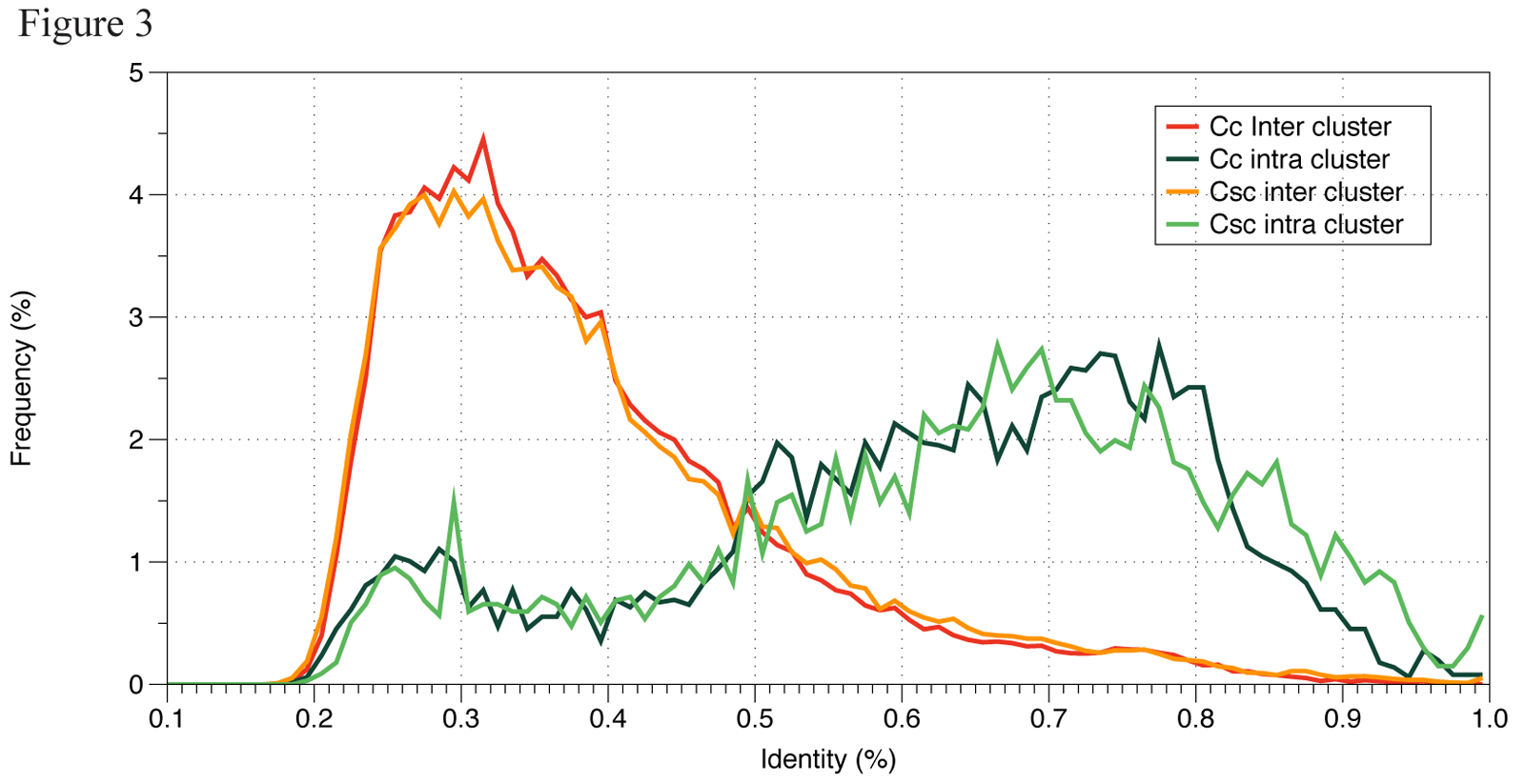
**Percentage identity distribution of NBS-encoding genes in *C. clementina* (Cc) and *C. sinensis* China (Csc).** Green lines indicate pairwise identity distribution of inter-cluster NBS-encoding genes. Orange lines indicate intro-cluster NBS-encoding genes.

We identified 254 and 246 gene conversion events from 76 NBS gene clusters in *C. clementina* and 75 NBS gene clusters in *C. sinensis* China, respectively (Table 4). The gene conversion events in *C. sinensis* USA is much less due to fragmental assembly. It is interesting that most of the conversion events (483 out of 520) were identified from the relatively small clusters with less than 10 NBS genes. Among these conversions, 119 events located in N-terminal, 178 in NBS domains and 223 in C-terminal, which indicating that there was no significantly bias in the location of conversion. Most gene conversions were between genes from the same group. We identified 101, 184 and 229 conversion events from NBS genes in TIR, CC1 and CC2 groups respectively and only four conversion events were identified between NBS genes of TIR and CC2 groups. While we could identify similar total numbers of conversion events in *C. clementina* and *C. sinensis* China, we found almost double such events in *C. sinensis* China TIR clusters versus in *C. clementina* TIR clusters. Meanwhile, we also found many more conversion events in *C. clementina* CC2 clusters than that in *C. sinensis* China CC2 clusters (Table 4).

**Table 4.** Gene conversion events found in *C. clementina* and *C. sinensis* China

| Citrus species | Cluster type | Number of clusters (with conversion events) | Number of conversion events | Affected genes | Mean size of conversion tracts |
| --- | --- | --- | --- | --- | --- |
| *Citrus clementina* | TIR | 27 | 35 | 32 | 111 |
|  | CC1 | 43 | 86 | 67 | 137 |
|  | CC2 | 46 | 133 | 83 | 146 |
| *Citrus sinensis* China | TIR | 36 | 69 | 42 | 137 |
|  | CC1 | 51 | 89 | 48 | 162 |
|  | CC2 | 57 | 92 | 75 | 116 |

### Analysis of NBS Orthologs

We identified 719 *Citrus* NBS gene pairs of orthologs amongst *Citrus clementina*, *C. sinensis* China and *C. sinensis* USA. 270 orthologs were shared between *C. clementina* and *C. sinensis* China (Figure S7); 227 orthologous gene pairs were shared between *C. clementina* and *C. sinensis* USA and 222 orthologous gene pairs were shared between *C. sinensis* China and *C. sinensis* USA. The percentages of identities between orthologous genes range from 53.8% to 100% (mean of 93.6%, median of 96.8%). The percentages of identities between orthologous genes from TIR group (TNL and TN, 88.42 ± 10.04) were significantly (T-test: P<2.6e-6) lower than those from CC groups (CNL and CN, 92.92 ± 7.03). The nonsynonymous divergence (dN) values of the 719 orthologs ranged from 1.46e-6 to 0.31 (mean of 0.03, median of 0.08) and the synonymous divergence (dS) values ranged from 1.14e-5 to 0.75 (mean of 0.05, median of 0.019) (Figure 4). The dN/dS ratios ranged from 0.001 to 50 with a median of 0.62. Similar to *Arabidopsis*, the dN and dS of orthologs from TIR group (TNL and TN) were higher than those from CC groups (CNL and CN). The dN values of orthologs from TIR group ranged from 0.001 to 0.24 (mean of 0.038, median of 0.017) while those of orthologs from CC group ranged from 1.4e-6 to 0.23 (mean of 0.025, median of 0.007). The dS of orthologs from TIR group ranged from 3.27e-5 to 0.76 (mean of 0.078, median of 0.029) while those of orthologs from CC group ranged from 1e-5 to 49 (mean of 0.04, median of 0.016). However, the dN/dS ratios of orthologs from TIR group were relatively lower than that of orthologs from CC group (Figure 4(A)).

**Figure 4.**
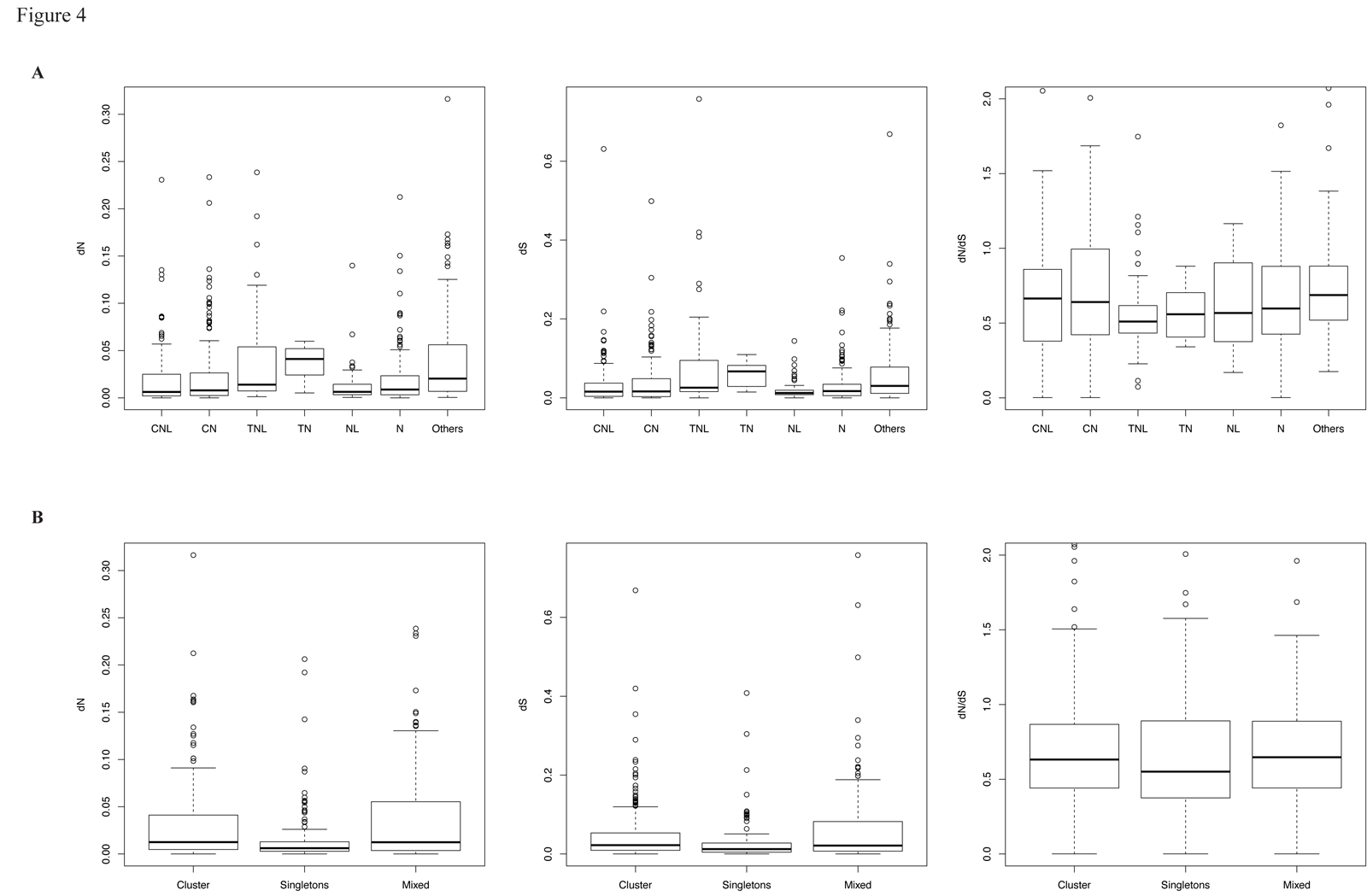
**Divergence of orthologous NBS-encoding genes amongst *Citrus* species.** A) Divergence of different classes. B. Divergence of singletons and cluster members.

The dN and dS rates as well as dN/dS rate ratios of ortholog residues in clusters are generally greater than those of orthologous singletons (Figure 4(B)). The median dN/dS ratio of 148 orthologous singletons was 0.55 while that of 361 orthologs in clusters was 0.63. There were 124 orthologs with dN/dS ratios above 1. These orthologs could undergo positive selection pressure. Most of the positive selected orthologs (87.9%, 109 out of 124) belonged to CC groups (49 orthologs from CC1 group and 50 from CC2 group).

We also detected 38 NBS gene syntenic blocks between *C. clementina* and *C. sinensis* China using MCScanX [29] (Figure S8). On average, there are 11 genes per block. The biggest block contains 45 NBS genes and was located in scaffold_5 of *C. clementina* and chromosome 3 of *C. sinensis* China. There were 80, 77 and 71 syntenic NBS genes in scaffold_5 of *C. clementina*/chromosome 5 of *C. sinensis* China, scaffold_3 of *C. clementina*/chromosome 5 of *C. sinensis* China and scaffold_7 of *C. clementina*/chromosome 1 of *C. sinensis* China. Furthermore, there were 68, 66 and 8 syntenic NBS genes in the unknown chromosome of *C. sinensis* China corresponding to NBS genes in scaffold three, five and seven of *C. clementina*, respectively.

### Analysis of Conserved NBS Gene Clusters

We identified 118 pairs of conserved NBS gene clusters between *C. clementina* and *C. sinensis* China (Figure S9). There were 19 NBS gene clusters completely conserved in *C. clementina* and *C. sinensis* China. For example, all 14 NBS genes in cluster CL225 of *C. sinensis* China have orthologs in the cluster CL168 *C. clementina* which contains nine NBS genes and vice versa. Some clusters (in both *C. clementina* and *C. sinensis* China) have several corresponding conserved clusters. This may due to either the genome arrangement or the incomplete assembly of genomes. Furthermore, there were 19 clusters of *C. clementina* with no conserved clusters in *C. sinensis* China and 18 clusters of *C. sinensis* China with no conserved clusters in *C. clementina*.

The conserved clusters provided additional insights into NBS gene evolution within and between *Citrus* genomes. For example, cluster CL142 (9 NBS genes) in *C. clementina* and cluster CL282 (7 NBS genes) in *C. sinensis* China were highly conserved. The phylogenetic tree of these 16 genes suggested division into two subgroups (depicted in blue and orange color in Figure 5(A)). This division indicates two ancestral genes for this conserved cluster. Using the phylogenetic tree as a framework, we reconstructed the evolutionary history of these two clusters. There were several tandem duplication events in the evolutionary history of the conserved clusters, and an extra tandem duplication was observed in *C. clementina* after it separated from *C. sinensis* (Figure 5(B)). Two NBS genes were lost in *C. clementina* and one NBS gene was lost in *C. sinensis*. Furthermore, there was a recombination event within *C. clementina*.

**Figure 5.**
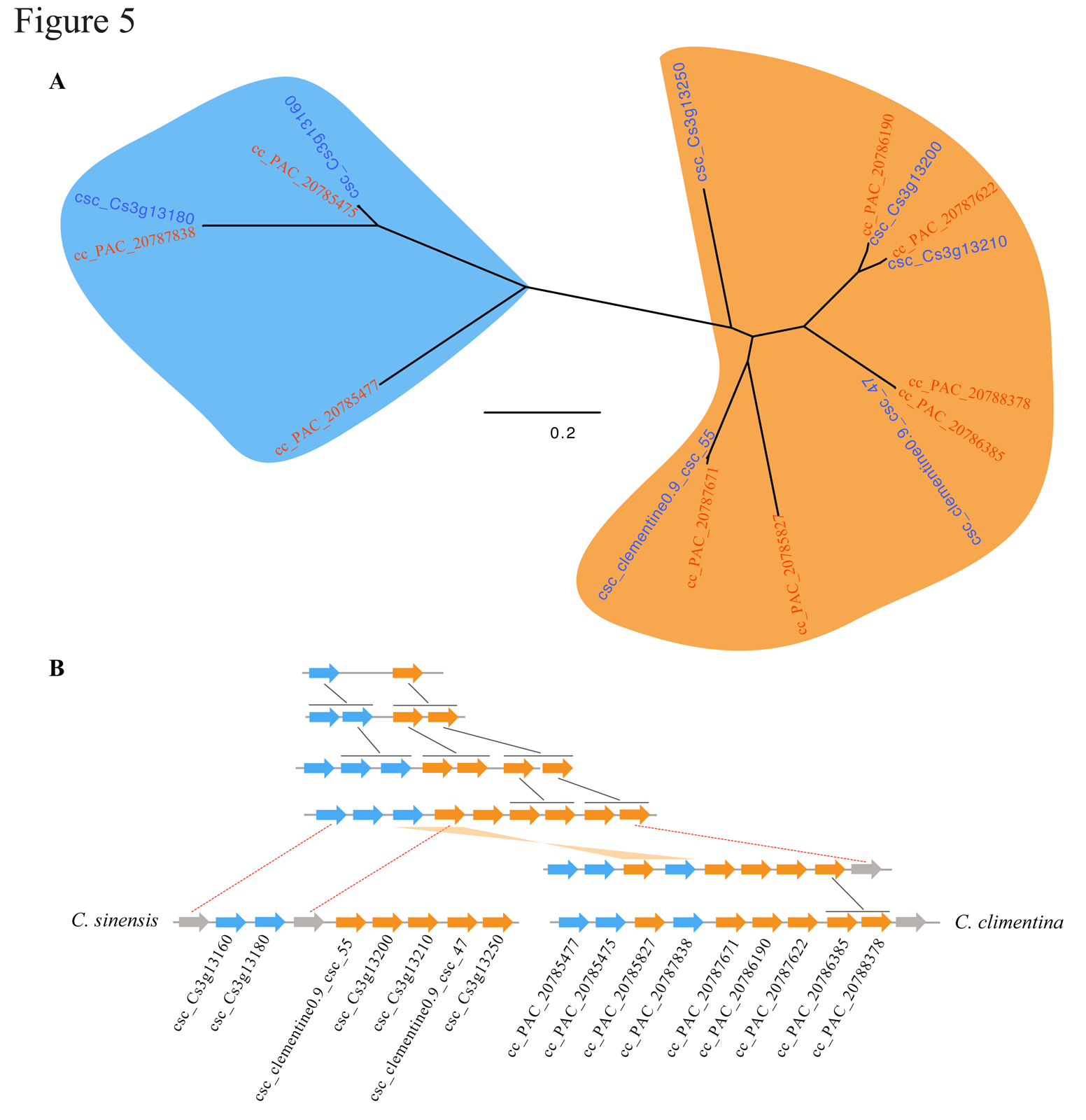
**Duplication histories of one of the conserved NBS-encoding genes cluster in *C. clementina* and *C. sinensis*. A**. Phylogenetic tree of the NBS-encoding genes; B. Orthologs in the conserved cluster of *C. clementina* and *C. sinensis*. Black connectors indicate tandem duplications and red dotted connectors indicate gene loss. A gene recombination event occurred in the ancestor of *C. clementina*, as indicated with an orange connector.

### Mutations and Transposons in *Citrus* NBS Genes

Plant NBS genes are continuously evolving. Sequence variation and structural constraints are shaped by gene birth-and-death processes [30, 31]. Besides possible interallelic recombination and gene conversion, gene mutations and transposable elements appear to play important roles in NBS gene evolution.

We compared the DNA sequences of NBS genes from different *Citrus* genomes to identify mutations. Approximately half of *Citrus* NBS genes have mutations maintained amongst corresponding orthologs. When a mutation took place in an exon and resulted in stop-codon gaining or frame-shift, the target gene often became a pseudogene. For example, Cs1g18610.1_cc_32, Cs1g18610.1 and orange1.1g003367m are orthologs of *C. clementina*, *C. sinensis* China and *C. sinensis* USA. There were 2 stop-codon gaining mutations in Cs1g18610.1, which means that Cs1g18610.1 might become a pseudo (nonfunctional) gene after gaining these mutations.

LTR retrotransposons are widespread in eukaryotic genomes, especially plant genomes. We predicted 19014, 6296 and 1479 LTR retrotransponsons with typical LTR characters in the draft genomes of *C. clementina*, *C. sinensis* China and *C. sinensis* USA, respectively. Then, we filtered out LTR retrotransponsons with low similarity with known TE proteins using BLASTX with e-value greater than 1e-5. Finally, 4920, 3726 and 1240 LTR retrotransponsons remained in *C. clementina*, *C. sinensis* China and *C. sinensis* USA, respectively. We identified 33, 32 and 4 NBS genes that were inserted with LTR retrotransposons in *C. clementina*, *C. sinensis* China and *C. sinensis* USA respectively (Table S6). Most of these genes will likely become pseudogenes due to these insertions. For an instance, orange1.1g043039m, orange1.1g043039m_cc_116 and orange1.1g043039m_csc_123 were orthologs of *C. sinensis* USA, *C. clementina* and *C. sinensis* China respectively (Figure S10(A)). The results of gene structure analysis showed that structure in orange1.1g043039m in *C. sinensis* USA seems relatively well maintained. But there is an about 10 kb fragment of LTR-retrotransposons in the corresponding homologous gene in *C. clementina* and *C. sinensis* China (Figure S10(B)). We identified both orange1.1g043039m_cc_116 and orange1.1g043039m_csc_123 as pseudogenes using PseudoPipe [32].

### Experimental Validation of One NBS Gene in *Citrus* Genomes

We validated the orthologs of a conserved NB gene, Cs1g09350.1, in a wide range of *Citrus* species. Cs1g09350.1 conserved in the 3 sequenced *Citrus* genomes. It is a CNL NBS gene and has 5 exons with about 4 kb in length. We designed the primers targeting about 3.5 kb fragment and could amplify a 3.5 kb fragment using PCR in different *Citrus* species, including *C. sinensis* (sweet orange), *C. clementina* (clementine), *C. japonica* (Kumquat), *C. sinensis* Navelina (Navel orange), *C. maxima* (pomelo), *C. aurantiifolia* (lime), and *C. aurantium* (sour orange). The orthologs from *C. sinensis* and *C. clementina* were identical as expected. There are a few mutations in the orthologs of *C. japonica*, *C. sinensis* Navelina (Navel orange), and *C. maxima*, but the orthologs of *C. aurantiifolia* and *C. aurantium* should be undergoing a pseudogenization process. There was a deletion plus several mutations in the NBS domain in *C. aurantiifolia*. There was an eight-base deletion in the LRR domain in *C. aurantiifolia* and *C. aurantium*. Most *Citrus aurantiifolia* mutations were shared with *C. aurantium* (Figure S11), and these mutations were not shared with other species. The results suggest that in *C. aurantiifolia* and *C. aurantium*, this gene is likely derived from the same common ancestor and was inherited as a pseudogene.

## Discussion

### Possible Common Ancestor of NBS Genes in Hybrid *Citrus sinensis* and Original *Citrus clementina* **Genome**

After carefully reannotating the *Citrus* genome sequences, we found similar numbers of NBS genes in *Citrus clementina* and *Citrus sinensis* China. There are slightly fewer NBS genes in *Citrus sinensis* USA, possibly due to the more fragmental assembly of this genome. In phylogenetic tree using NBS domains, the NBS genes from three different *Citrus* genomes are mixed together. After partitioning NBS genes on the tree into clades, we found that 97.7% of the clades containing more than three genes had members from each of three *Citrus* genomes. This pattern suggests that these three *Citrus* genomes have similar NBS genes derived from common ancestors. Because *Citrus sinensis* is the hybrid of *C. reticulata* (Mandarin orange) and *C. maxima* (pomelo), it should be heterozygous and some of NBS genes in *Citrus sinensis* are expected have different genetic distances from NBS genes in *Citrus clementine.* This would support previous observations of NBS genes in F1 progeny of *Poncirus trifoliata* (trifoliate orange) and *Citrus reticulata* (tangerine). However, this is clearly not the case.

Furthermore, we mapped the resequenced reads of three *C. maxima* (pomelo) genomes and three *C. reticulata* (Mandarin orange) genomes onto the genome of *Citrus sinensis* China. In this case, 62% of *C. sinensis* China NBS genes have a copy present in all six resequenced genomes and 99% of *C. sinensis* China NBS genes have a copy in at least one of the resequenced genomes. The mapping results confirmed that a significant percentage of NBS genes of hybrid *C. sinensis* genomes have corresponding homologous genes in both the *C. maxima* and the *C. reticulata* genomes. Because the reference genome sequence of *C. maxima* is not yet available, the total number of NBS genes in *C. maxima* genomes is still not known. However, we can at least conclude from the mapping of resequenced genomes that the *C. maxima* genome has homologous copies of NBS genes in *C. reticulata* and *C*. *sinensis* genomes. The homologous NBS genes in *C. maxima* and *C. reticulata* may be the reason that NBS genes in their hybrid *Citrus sinensis* are similar to those in *C. reticulata* in this study.

### Three Groups of *Citrus* NBS Genes

We identified 442, 393 and 264 genes with full length NBS domains from *Citrus clementina*, *Citrus sinensis* China and *Citrus sinensis* USA reference genomes, respectively. There are also many genes with short NBS domains in three *Citrus* genomes. The *Citrus* NBS genes can be divided into three groups according to the phylogenetic tree of NBS domains: two of them contain CC domain and the other group contains TIR domain. The number of CC NBS genes is three times of the number of TIR NBS genes. In most of the TIR NBS genes, we can find the LRR domains defined in Pfam database [33]. We only can identify LRR domains in small part of CC NBS genes using Pfam LRR domain definition. However, we can find the LxxLs repeats in most of the CC NBS genes as shown in motifs from MEME. This implied that there may be other types of LRR domain in *Citrus* CC NBS genes. Our motif analyses also showed that motifs of TIR domains in *Citrus* CC2 and TIR NBS genes are similar to those of TIR domains in *Arabidopsis* [23] and *Populus trichocarpa* [24] TIR NBS genes. Furthermore, the motifs of *Citrus* CC domains in CC1 NBS genes are similar to those motifs of CC domains in Arabidopsis CC NBS genes and the motifs of *Citrus* CC domains in *Citrus* CC2 NBS genes are similar to those motifs of CC domains in japonica rice CC NBS genes [28]. The different structure of motifs in NBS domain and CC domain between *Citrus* CC1 NBS genes and *Citrus* CC2 NBS genes implied that they are from different evolutionary origin.

To further confirm the three groups of NBS genes, we identified the NBS genes from *Arabidopsis*, *Populus*, *Oryza sativa* and grape. We used the same criteria to select the NBS domains from NBS genes of these genomes. Finally, we selected 152, 216, 209, 126 NBS domains from *Arabidopsis*, *Populus*, *Oryza sativa* and grape, respectively. Then, together with 442 NBS domains selected from *Citrus clementina,* we constructed a phylogenetic tree of those 1145 NBS domain sequences from five genomes. As shown in Figure 6, the un-rooted phylogenetic tree was divided into three main branches. The *Populus,* grape and *Citrus* genomes have significant amount of NBS genes in all three branches. However, the NBS genes of *Oryza sativa* dominated in CC2 branch and most of *Arabidopsis* NBS genes located in CC1 and TIR branches. Our study showed that the NBS genes can be divided into three major groups as the NBS genes with the CC domains are separated into two groups. The three groups of NBS genes underwent divergent evolution in different genomes. Further comparison of NBS genes of more genomes may help to understand the evolution of NBS genes and will help elucidate how plants maintain and adapt their defense system against pathogens.

**Figure 6.**
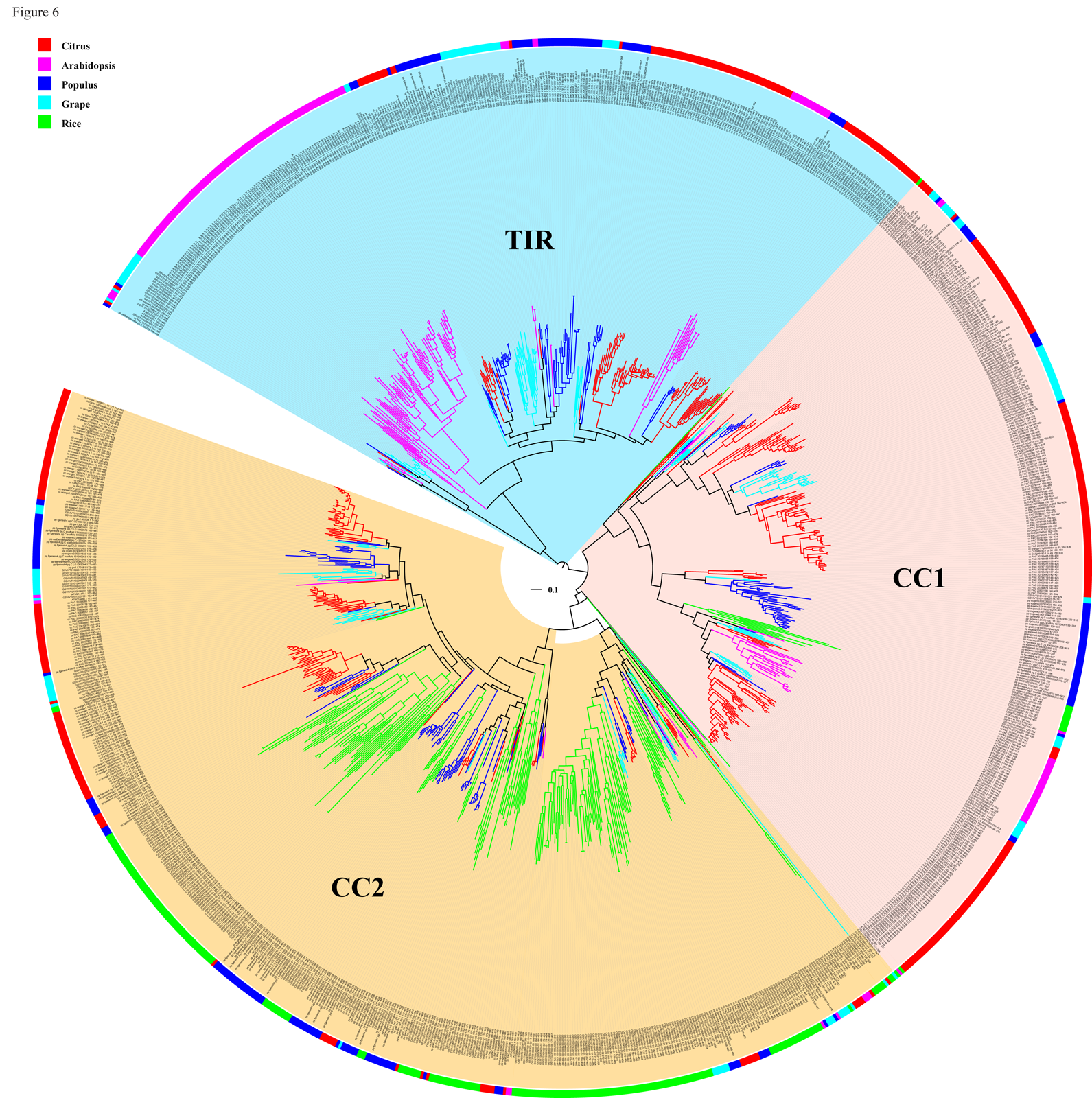
**Phylogenetic analysis of NBS genes of *Citrus clementina*, *Arabidopsis thaliana, Populus trichocarpa, Oryza sativa* and *Vitis (grape)***. Species are indicated by the color of the outer circle and branches.

### Highly Clustering of *Citrus* NBS Genes

The *Citrus* NBS genes are highly clustered in the genome. 84.9% of NBS genes in *Citrus clementina* and 76.9% of NBS genes in *Citrus sinensis* China were found in clusters. Previous studies showed 76% of rice [34], 64% of *A. lyrata* and 71% *A. thaliana* [15], 83.2% of grapevine and 67.5% of poplar [11] NBS genes are found in clusters. The percentage of *Citrus* NBS genes in clusters is in the high level comparing to other genomes. The average number of genes per clusters is 4.86 in *Citrus clementina* and 3.97 in *Citrus sinensis* China. These numbers are similar to those in other genomes [11]. Furthermore, most *Citrus* NBS genes in the same cluster belong to the same phylogenetic group, suggesting that tandem duplication is the primary mechanism for the expansion of NBS genes in the *Citrus* genus.

### Molecular Evolution of *Citrus* NBS Genes

Similar to NBS genes in other genomes, *Citrus* NBS genes are highly dynamic and are shaped by several evolutionary processes leading to several differences amongst NBS genes, including domain presence and mutation constraints and genome organization. Our results revealed multiple molecular evolution events amongst *Citrus* NBS genes including gene duplications, gene conversions, mutation constraint changes, recombination and transposable element insertions. Likely these events support a birth-and-death process leading to the origins of new NBS genes as well as malfunction and loss of other NBS genes [14]. We found more than 200 tandem duplications in both *C. clementina* and *C. sinensis* China genomes alone. We also found that NBS genes become pseudogenes following original frame-shift mutations leading to mutation accumulations plus disruption of gene constraint leading to loss of function through transposable element insertions.

Most molecular evolutionary events occurred within NBS gene groups, suggesting that gene birth-death processes have been occurring since divergence from a common ancestral gene copy. Interestingly, the molecular evolution processes occurred differently amongst NBS gene groups and differently within each *Citrus* genome. For example, the number of gene conversion events in *C. sinensis* China TIR clusters is almost double that in *C. clementina* TIR clusters, while there are many more conversion events in *C. clementina* CC2 clusters than in *C. sinensis* China CC2 clusters (Table 4). Numbers of tandem duplications in CC1 and CC2 groups are much greater than that from the TIR group.

## Conclusions

Our comparative analyses yield valuable insight into the understanding of the structure, evolution and organization of NBS genes in *Citrus* genomes. There are significantly more NBS genes in *Citrus* genomes compared to other plant species. *Citrus* NBS genes are structurally highly clustered. NBS genes in hybrid *C. sinensis* genomes are very similar to those in progenitor *C. clementina* genomes and their NBS genes may be derived from possible common ancestral gene copies. Furthermore, our comprehensive analysis also showed that there are three groups of plant NBS genes while NBS genes containing CC domains can be divided into two groups.

## Materials and Methods

### Sequences Used

We downloaded the draft genome sequences and the original gene annotations of *Citrus clementina* (clementine) and *Citrus sinensis* USA (sweet orange from USA) from the Citrus Genome Database (http://www.citrusgenomedb.org/) and those of *Citrus sinensis* China (Chinese sweet orange [21]) from the *Citrus sinensis* annotation project (http://citrus.hzau.edu.cn/orange/). The sizes of assembled genomes of *Citrus clementina*, *Citrus sinensis* China and *Citrus sinensis* USA are 301, 328 and 319 million basepairs, respectively. The *C. clementina* genomes assembled into nine major scaffolds and 95.8% of the sequences were assigned to those nine scaffolds. About 72.9% of the *C. sinensis* China genome assigned to nine chromosomes. The genome of *C. sinensis* USA only assembled to 12,574 scaffolds and the N50 of scaffolds is 250 kb. The original gene annotations of *Citrus clementina*, *Citrus sinensis* China and *Citrus sinensis* USA have 24,533, 29,385 and 25,397 genes respectively. We also downloaded the resequence data of three *Citrus clementina* (clementine) and three *Citrus maxima* (pomelo) genomes from the *Citrus sinensis* annotation project (http://citrus.hzau.edu.cn/orange/).

### Identification of NBS Genes in *Citrus* Genomes

We first screened the original predicted citrus open reading frames (ORFs) using hmmsearch [35] with the hidden Markov models (HMM) of Pfam [36] for NBS domain presence (NB-ARC, PF00931) using an e-value cut-off of 0.1 for the hmmsearch. Then, the proteins selected by the HMM were searched against the Swiss protein database [37] to confirm the annotation using BLASTP [38]. Only the proteins that have a significant match (e-value < 1E-5 in BLASTP search) with the NBS proteins or resistant proteins in the Swiss protein database were identified as NBS-containing proteins. To recover possible NBS genes that may be missed in the original gene annotations, we mapped the identified NBS genes to the draft genome using TBLASTN. The matched sequences with e-value < 1E-5 were then predicted using Genewise [39]. The new genes predicted by Genewise were also confirmed using NBS domain HMM screening and by BLASTP searching through the Swiss protein database.

### Identification of Orthologous NBS genes in *C. clementina* and *C. sinensis*

Orthologous NBS genes of *C. clementina* and *C. sinensis* were identified using the reciprocal best blast method [40]. We used NBS genes of *C. clementina* as query sequences to search against the NBS genes of *C. sinensis* and *vice versa*. Protein pairs with reciprocal best hits of e-value < 1E-20 were defined as orthologs.

To calculate the rates of nonsynonymous, synonymous and their rate ratio (dN, dS and dN/dS) of orthologous pairs, we first aligned orthologous protein sequence pairs using mafft, and then converted the protein alignments to codon-based alignments using PAL2NAL [41]. We calculated the dN, dS and dN/dS ratios using the codeml program in PAML version 4.7 [42].

### Phylogenetic Analysis

We constructed the phylogenetic tree of NBS genes of the three citrus genomes using only the conserved NBS domain. Only 1,099 NBS domain sequences that have both the P-loop and the MHDV motifs and were longer than 70% (200 amino acids) of the full-length NBS domain were included. Next, NBS domain sequences were aligned in mafft [43] using an auto alignment model and a best fit maximum likelihood phylogenetic tree of NBS genes was constructed using FastTree [25] with default parameters (JTT+CAT). The resultant best fit phylogenetic tree was divided into 3 main groups based upon clade support. For each group, we constructed a new ML tree with *Streptomyces coelicolor* protein P25941 as the outgroup using FastTree [25]. The average identity of each group was calculated using alistat implemented in SQUID (http://selab.janelia.org/software.html). We further partitioned the NBS gene tree into clades using the depth-first phylogeny partition method in PhyloPart [44] with distance threshold 0.025. This clustered the NBS genes into 114 clades, which contain more than 2 genes, and 8 orphan genes.

### Domain and Motif Annotation

The Toll-Interleukin receptor (TIR) domains in *Citrus* NBS-containing proteins were identified using hmmsearch [35] with the HMM model of Pfam domain PF01582 and a 0.1 e-value cut-off. The Leucine-Rich Repeat (LRR) domains in *Citrus* NBS containing proteins were identified using the HMM models of Pfam LRR domains with e-value cut-off of 0.1. As long as there is a significant hit to one of LRR domain models (e-value < 0.1), it was define as an LRR-containing protein. We used MARCOIL [45] with a threshold probability of 90 and COILS [46] with a threshold of 0.9 to search for Coiled Coil (CC) domains in the N-terminal region of *Citrus* NBS-containing protein. We considered a protein as a CC-containing protein if either MARCOIL or COILS reported a CC domain in it.

We identified 20 motifs amongst the NBS genes in each of the three main phylogenetic groups separately using MEME SUITE [47]. The motif width was set to between 6 and 50 for MEME. Then, we searched the motif structure of all genes in each group using MAST with default parameters (-ev 10 −mt 0.0001).

### Pseudogene Identification

We identified possible pseudogenes using PseudoPipe [32] with default parameters (-e 0.1). The PseudoPipe algorithm identifies pseudo genes by integrating sequence similarity, intron-exon structure, plus presence of stop codons and frame-shifts. We used all *Citrus* NBS genes to search *Citrus* genomes for potential NGS pseudo genes.

### Transposon Identification

All long terminal repeat (LTR) retrotransponsons in each citrus genome were identified using LTR finder [48] with default parameters (-o 3 -t 1 −e 1 -m 2 -u -2). Then, we used a script program to match the location of the LTR transposons to the NBS-LRR genes in each citrus genome.

### NBS Gene Synteny Identification

Gene synteny was identified and defined using MCScanX [29] with default parameters (-A -u 5000). First, NBS genes between two genomes were aligned using BLASTP and the matches with E-value < 1e-5 were sorted according to their chromosome positions. Synteny scores were then calculated for each block based upon gene position. Two genes were considered in the same block if there were fewer than 25 genes separating them. MCScanX reported blocks with at least 5 collinear gene pairs.

### Gene Cluster Analysis

We grouped the NBS genes in each *Citrus* genome into the same cluster if the genome location between two genes was within 200 kb. We also identified the conserved gene clusters between the *C. clementina* and *C. sinensis*. If all genes in a cluster of *C. clementina* have orthologs in the corresponding cluster of *C. sinensis* and vice versa, then these two clusters were called completely conserved clusters. If only part of genes in the clusters have orthologs, we called these two clusters partially conserved.

### Gene Conversion Detection

We first aligned the sequences of NBS genes in the same cluster using mafft [43]. Then, we used GENECONV [49] version 1.81a with default settings (N=10,000) to detect gene conversions. GENECONV identifies gene conversions by finding identical fragments between pairs of sequences in a nucleotide alignment. A global *P* value ≤ 0.05 was used to assess the statistical significance of the observed conversions. GENECONV requires at least three sequences for analyses in order to account for shared ancestral states. Thus, we only detected conversions in clusters containing three or more genes.

### Tests for Sites under Positive Selection

The amino acid sequences of NBS genes from the same clade were aligned with mafft [43]. Then we converted the protein alignments to codon-based alignment using PAL2NAL [41]. The positively selected sites were statistically identified using the Bayesian approach implemented in codeml within PAML [42]. We also further examined sites in the ω > 1 class with >90% posterior probability.

### Mapping of Re-sequencing Data of *Citrus clementina* and *Citrus maxima*

We mapped the raw reads of each re-sequenced sample to the draft genome of *Citrus sinensis* China using BWA [27] with default parameters (-k 19 -d 100 -A 1 -B 4 -O 6). Then, the mapped reads of NBS genes regions were extracted using BEDtools [50]. Two types of coverage of each NBS gene in *Citrus sinensis* were calculated. One coverage type divides the length of mapped sequences by the whole gene sequence length and the other type divides the length of mapped sequences by the length of exon sequence only.

### *Citrus* DNA Extraction and PCR Amplication

*Citrus* leaf samples were collected from the six *Citrus* plants in USHRL's (USDA Horticultural Research Laboratory, Fort Pierce, Florida): *Citrus sinensis* (sweet, Navel orange), *Citrus aurantium* (Karum jamir, sour orange), *Citrus reticulata* (Mandarin orange), *Citrus clementina* (Clementine), *Citrus aurantiifolia* (sweet lime), *Citrus japonica* (Yuzu, kumquat), and *Citrus maxima* (pomelo). Total DNA was extracted from leaf midribs following the DNeasy® Plant Mini Kit standard protocol (Qiagen Inc., Valencia, CA), followed by DNA quantity and quality evaluation with Nanodrop. We chose the NBS gene, Cs1g09350, which was conserved in *C. clementina* and *C. sinensis* for validating the conservation of NBS gene among different *Citrus* genomes. Primers used in this study were designed using Oligo 7.23 (Molecular Biology Insights, Inc., Cascade, CO, USA). High Fidelity Platinum® Taq DNA Polymerase (Invitrogen, Carlsbad, CA, USA) was used to amplify the NBS-LRR genes from *Citrus* DNA. For PCR, 20 µL reactions using standard conditions provided by the manufacturer for High Fidelity Platinum® Taq DNA Polymerase. PCR was performed using an initial denaturation at 95°C for 3 minutes, 35 cycles of 94^o^C for 20 seconds, 50-52^o^C for 20 seconds (specified by different primer sets) and 68^o^C for 3 minutes, follow by final extension at 68^o^C for 10 minutes in a C1000^TM^ Thermal Cycler (Bio-Rad, Hercules, CA). The cloning and sequencing analysis of amplified PCR products were conducted as previously described [51]

## Authors’ Contribution

FL, YD, PS and LD designed the research. YW performed the bioinformatics studies. LZ performed the experimental verification. YW, DL, ALR and FL performed the genetic and evolution analyses. ALR, PS, LD, YD and FL wrote the manuscript. All authors read and approved the final manuscript

## Acknowledgement

This work was supported in part by the *Citrus* Research and Development Foundation, Inc.

---

**Figure S1.**
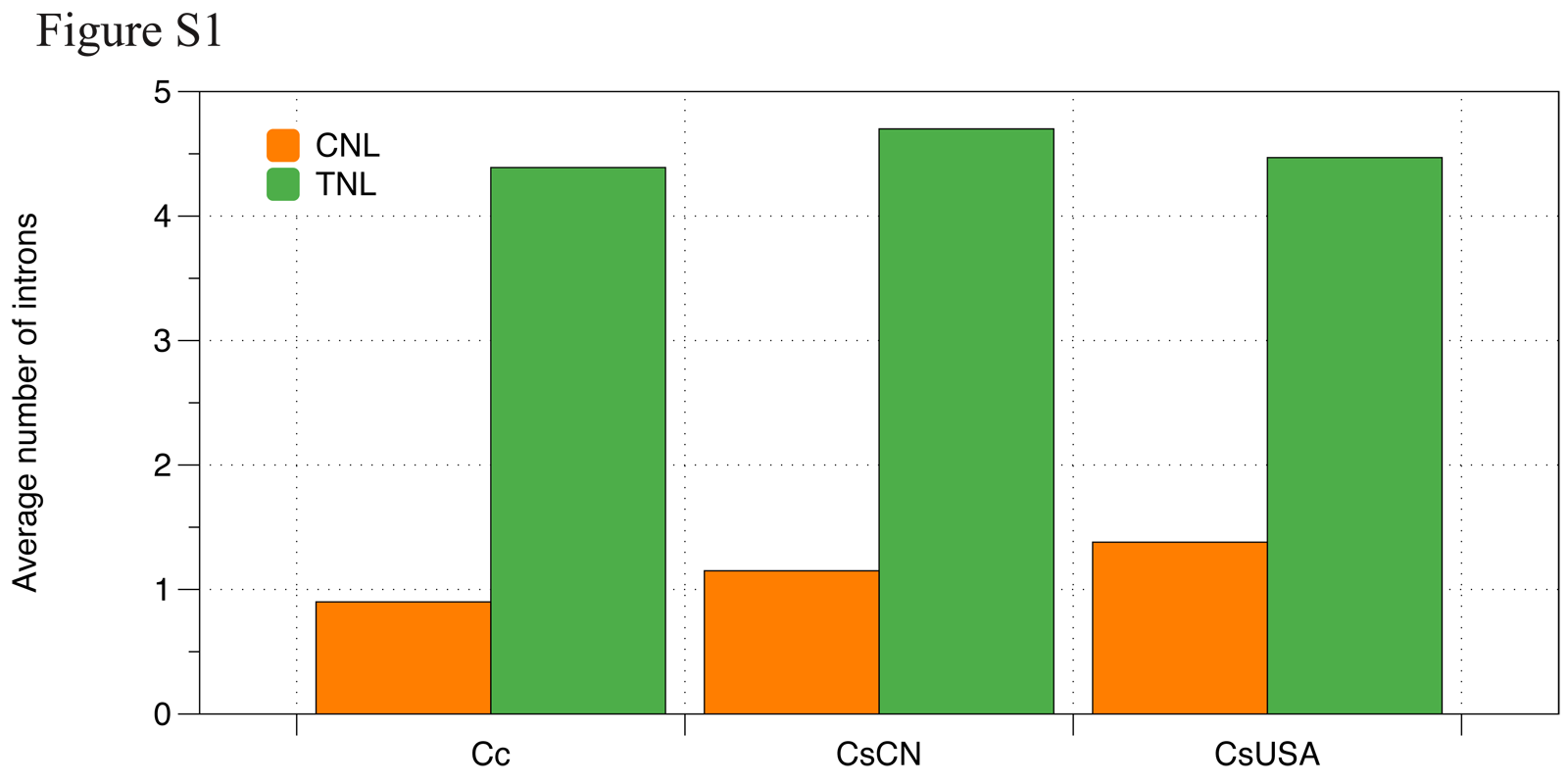
Average intron number of CNL and TNL. Cc: *C. clementina*, CsCN: *C. sinensis* China and CsUSA: *C. sinensis* USA.

**Figure S2.**
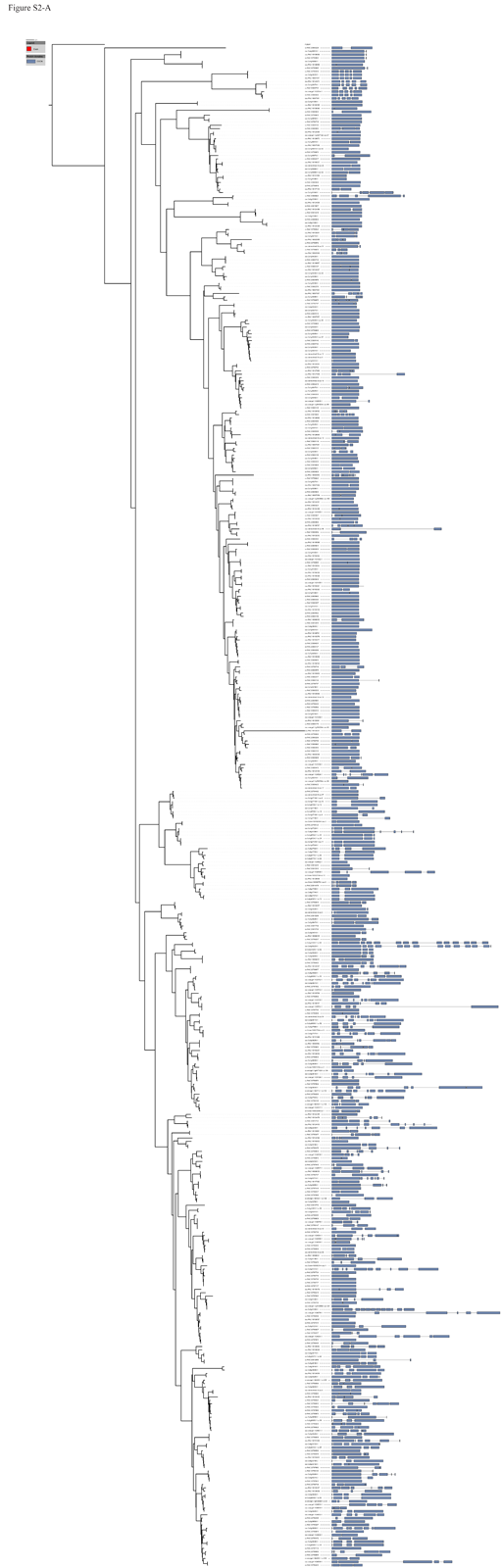
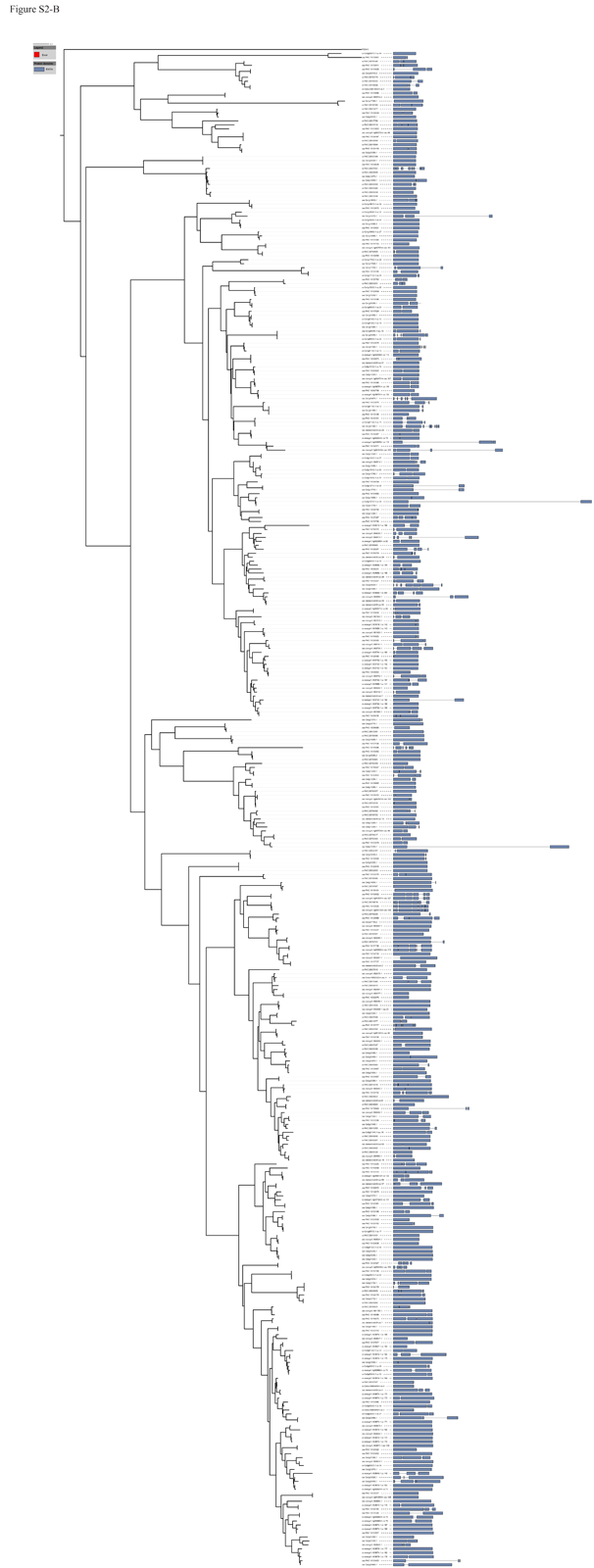
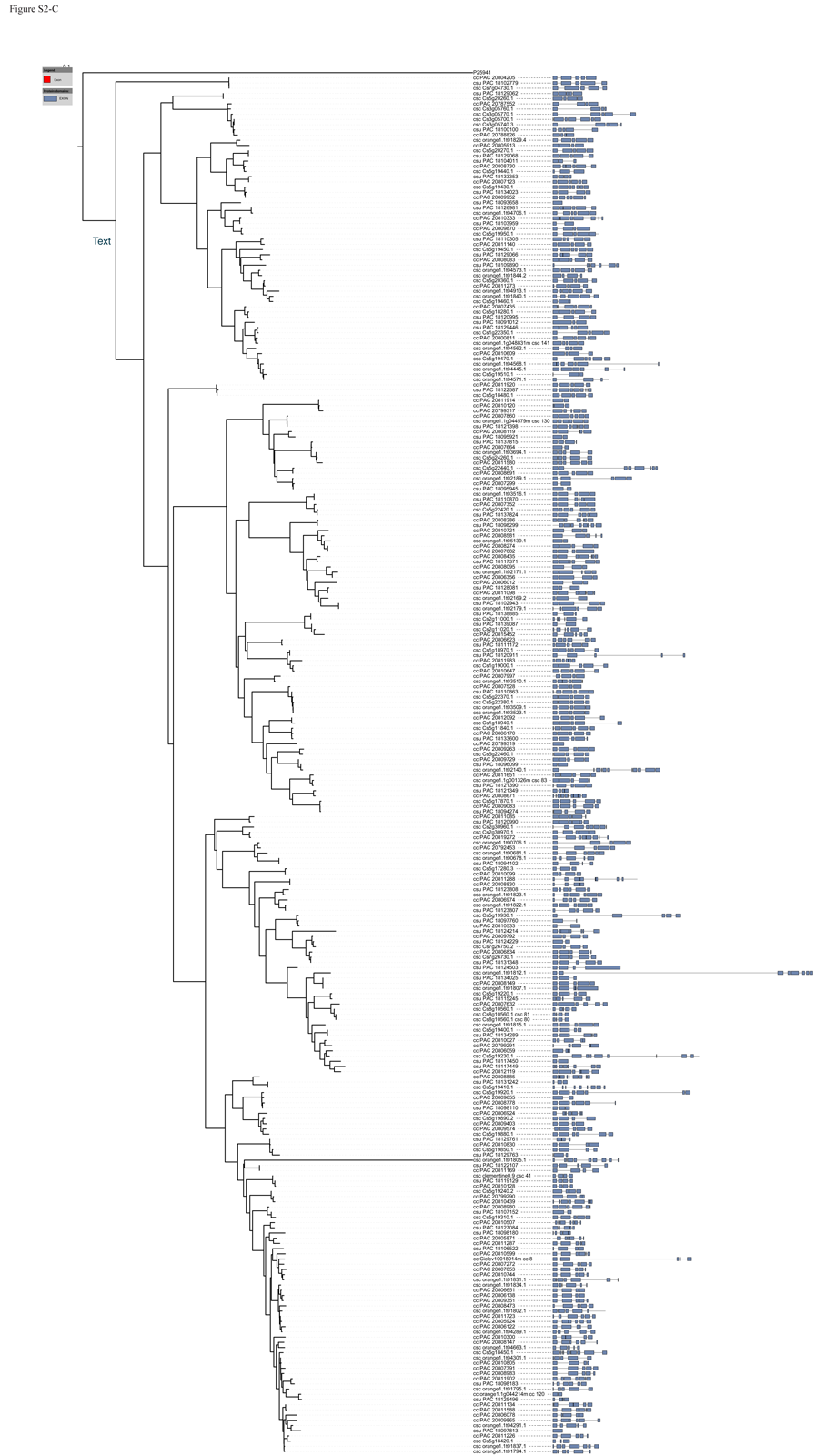
Phylogenetic trees of three groups of citrus NBS gene with P25941 as outgroup. A, CC1 group; B, CC2 group and C, TIR group.

**Figure S3.**
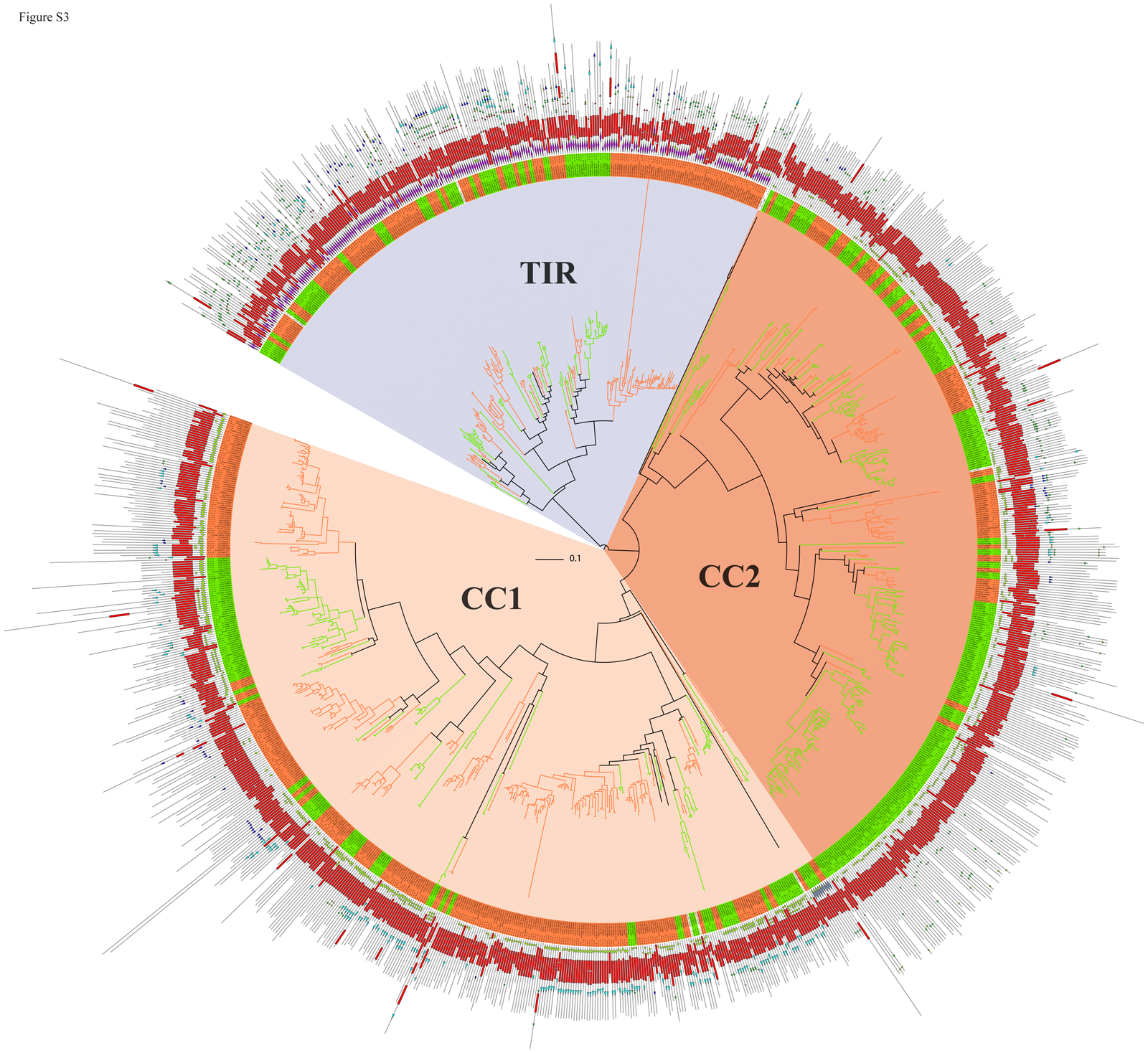
Structure of domains of citrus NBS genes. The red rectangles indicated the NBS domains, the green ellipses indicated CC domains, the left pointing pentagrams indicated TIR domains and the left pointing triangle indicated LRR domains.

**Figure S4.**
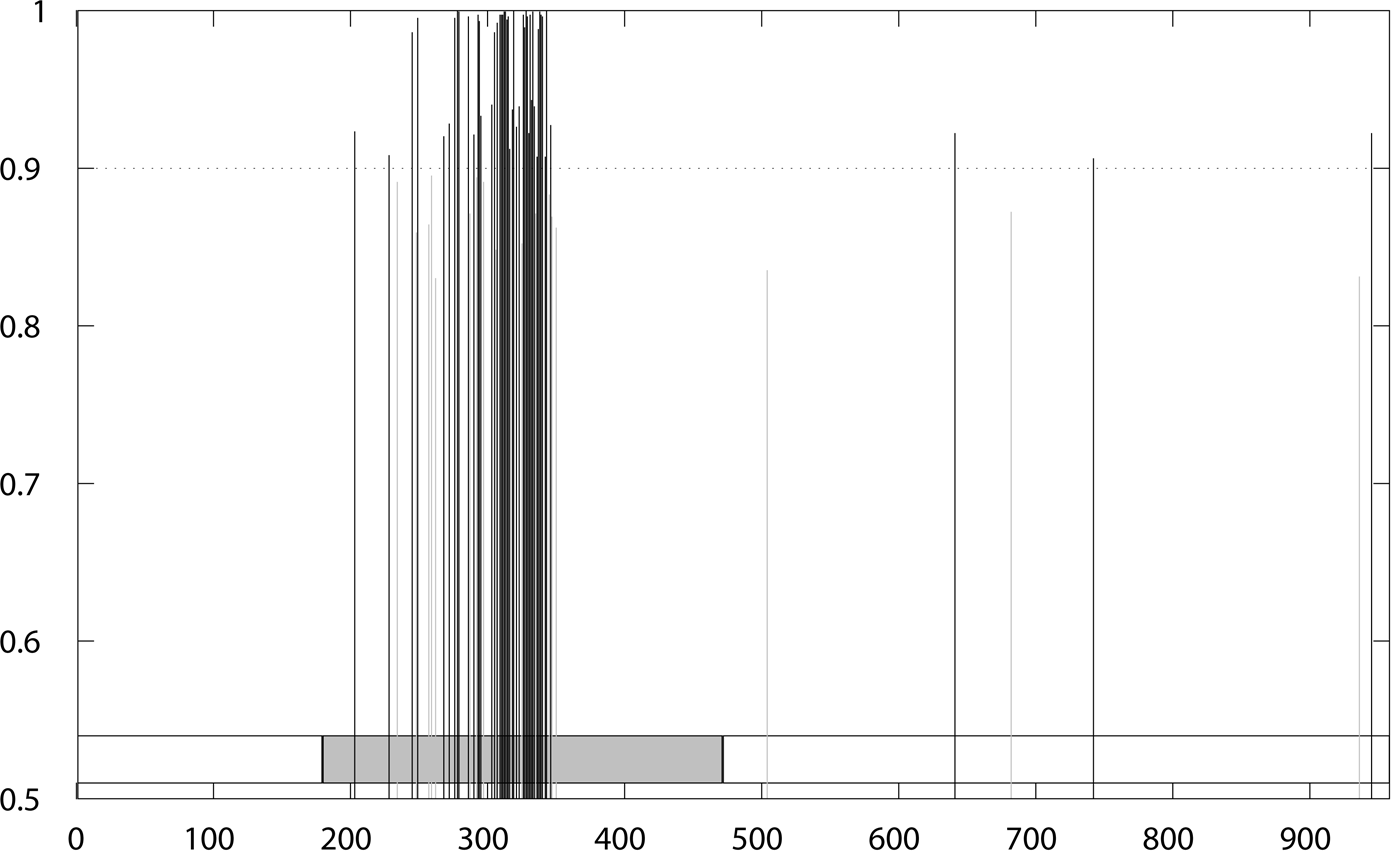
Positive selection sites of Clade_1260. The grey box indicated the NBS domain.

**Figure S5.**
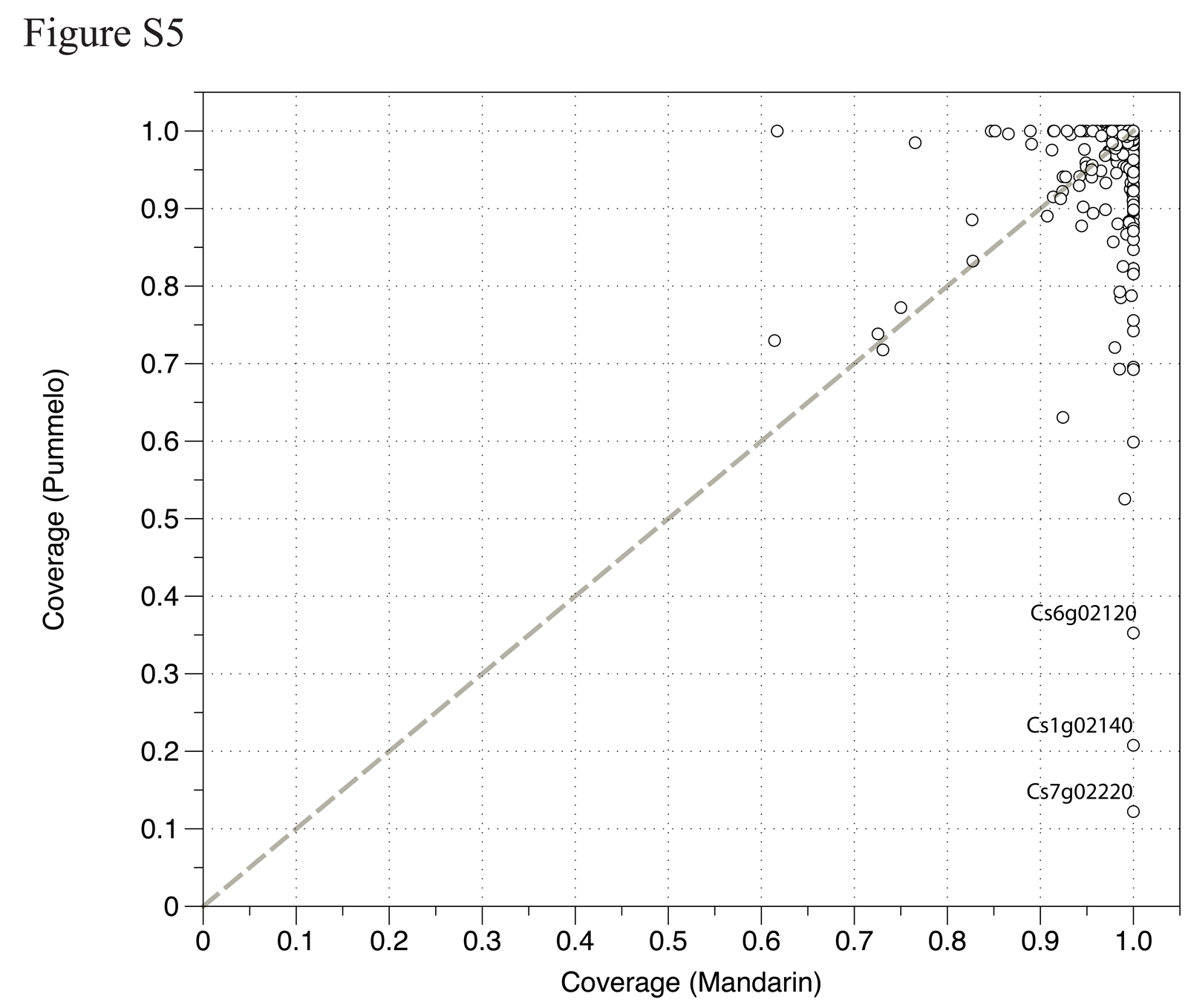
The exon coverage of NBS-encoding genes in the resequences of citrus genome (3 mandarin and 3 pomelo).

**Figure S6.**
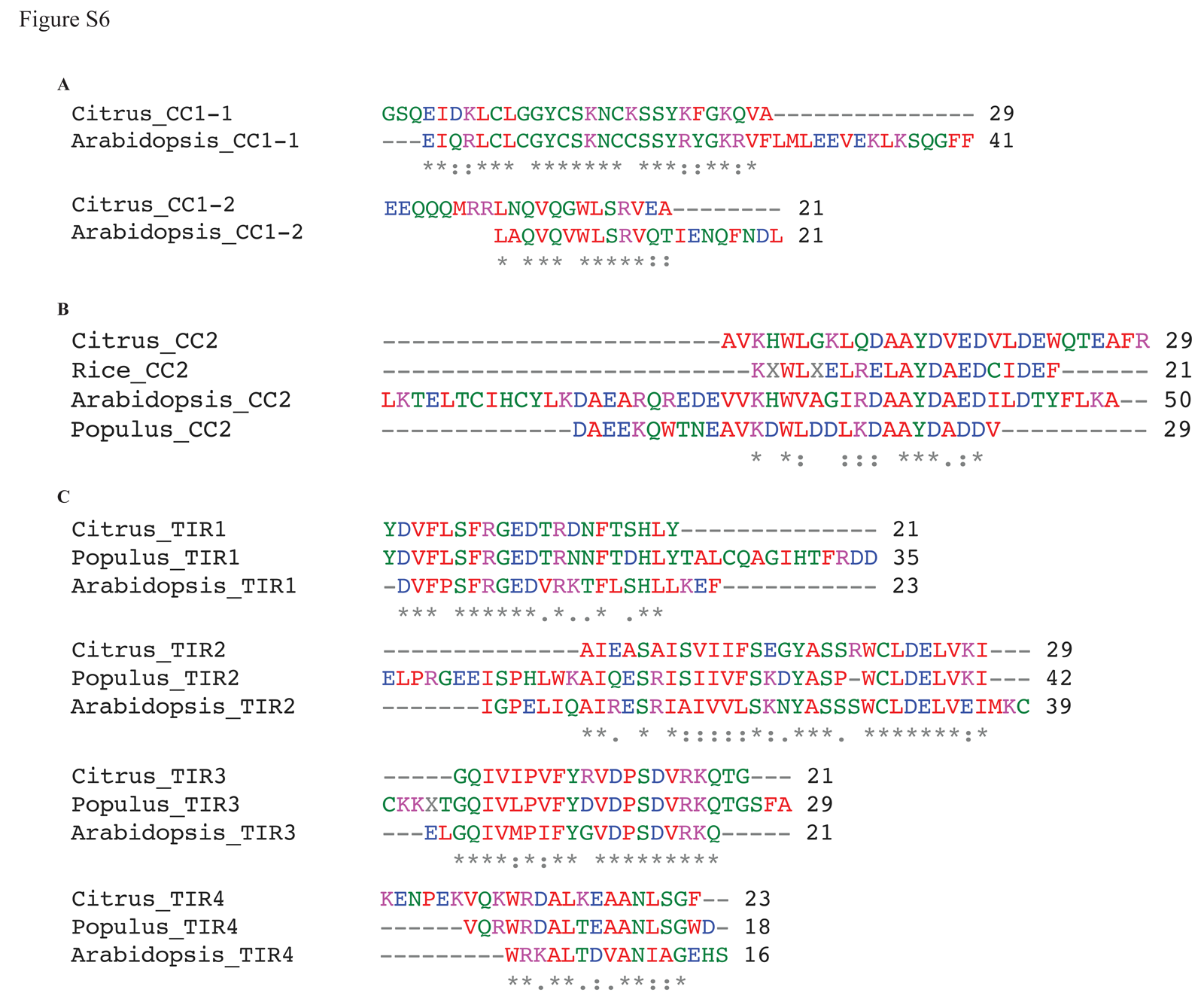
Multiple alignments of motifs from CC1 domain (A), CC2 domain (B) and TIR domain (C).

**Figure S7.**
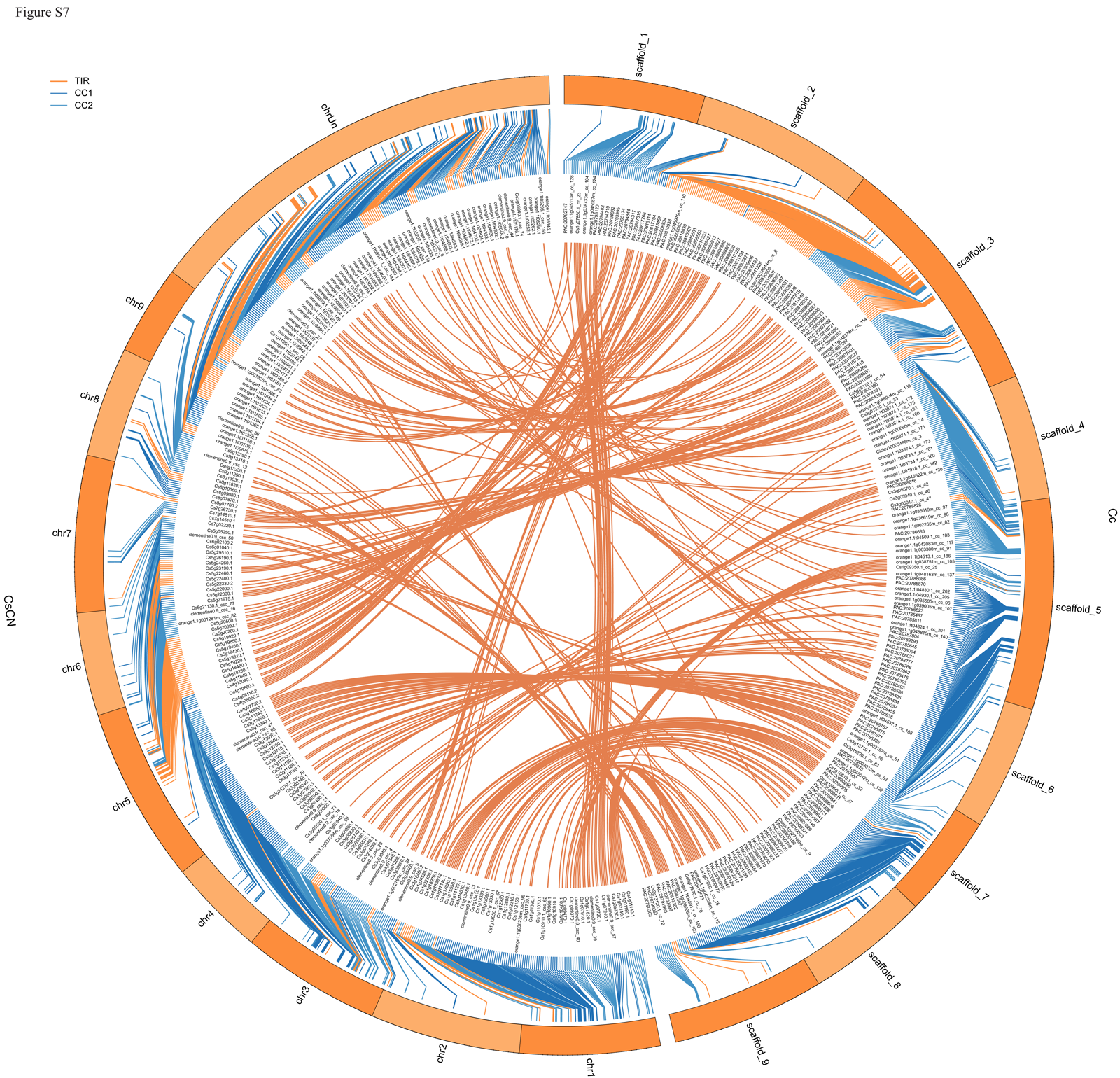
Orthologs of citrus NBS-encoding genes of Cc and CsCN and their genome locations. The outer blue circles indicated 9 largest scaffolds of Cc and orange ones indicated 9 chromosomes plus an un-located pseudo-chromosome. The NBS-encoding genes were arranged due to their locations on the chromosomes. The NBS genes in the TIR group were indicated with orange links, CC1 in blue and CC2 in light blue ones. The NBS-encoding orthologs of *C. clementina* and *C. sinensis* were indicated with the link lines.

**Figure S8.**
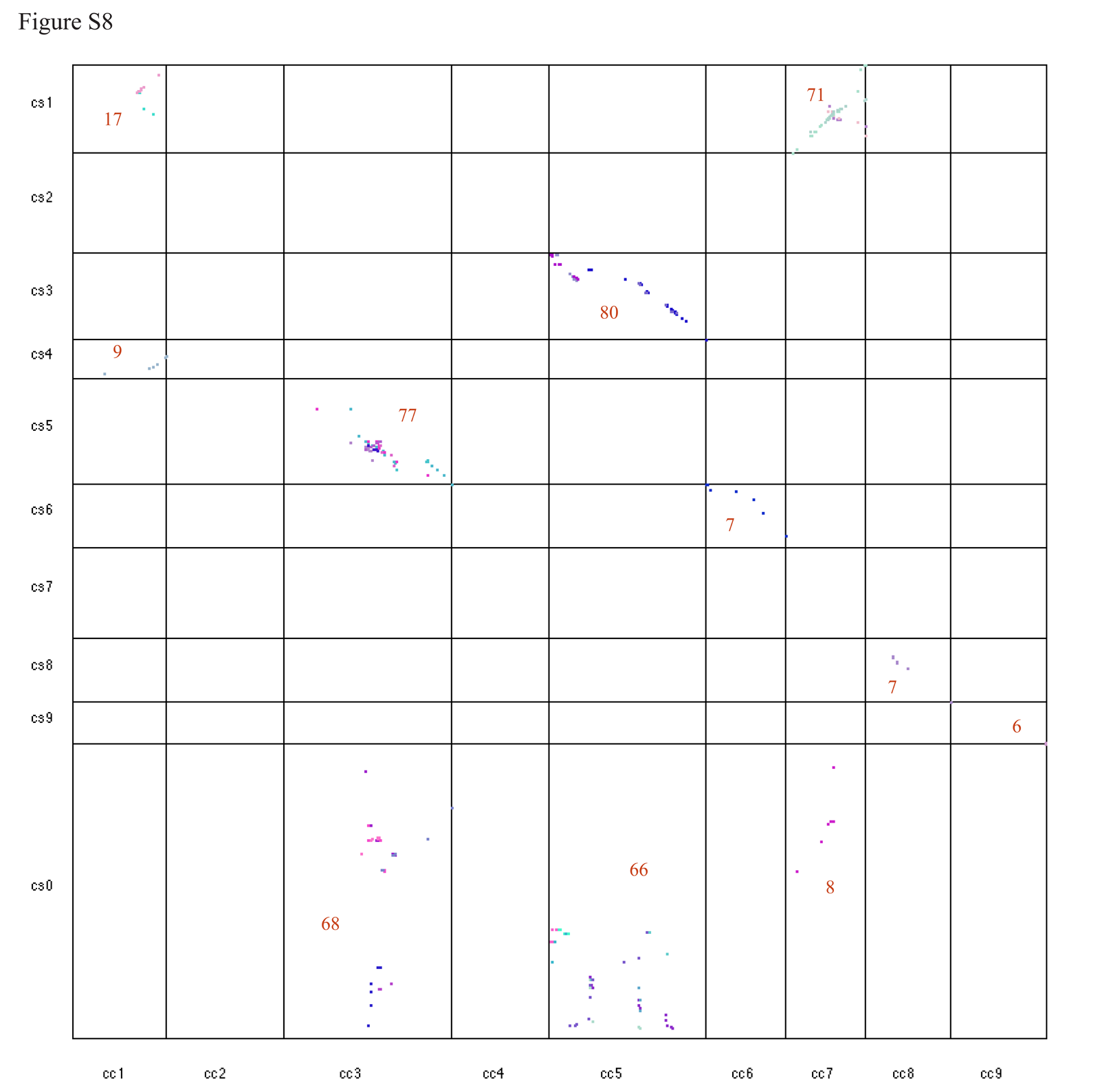
Dot plotting of syntenic orthologs of the NBS genes between *C. clementina* (Cc) and *C. sinensis* China (CsCN) produced by MCScanX. The numbers in the grids indicated the number of syntenic ortholog pairs in the corresponding scaffolds. The scaffolds of *C. clementina* (Cc) were arranged in x-axis and that of *C. sinensis* China (CsCN) were arranged in y-axis.

**Figure S9.**
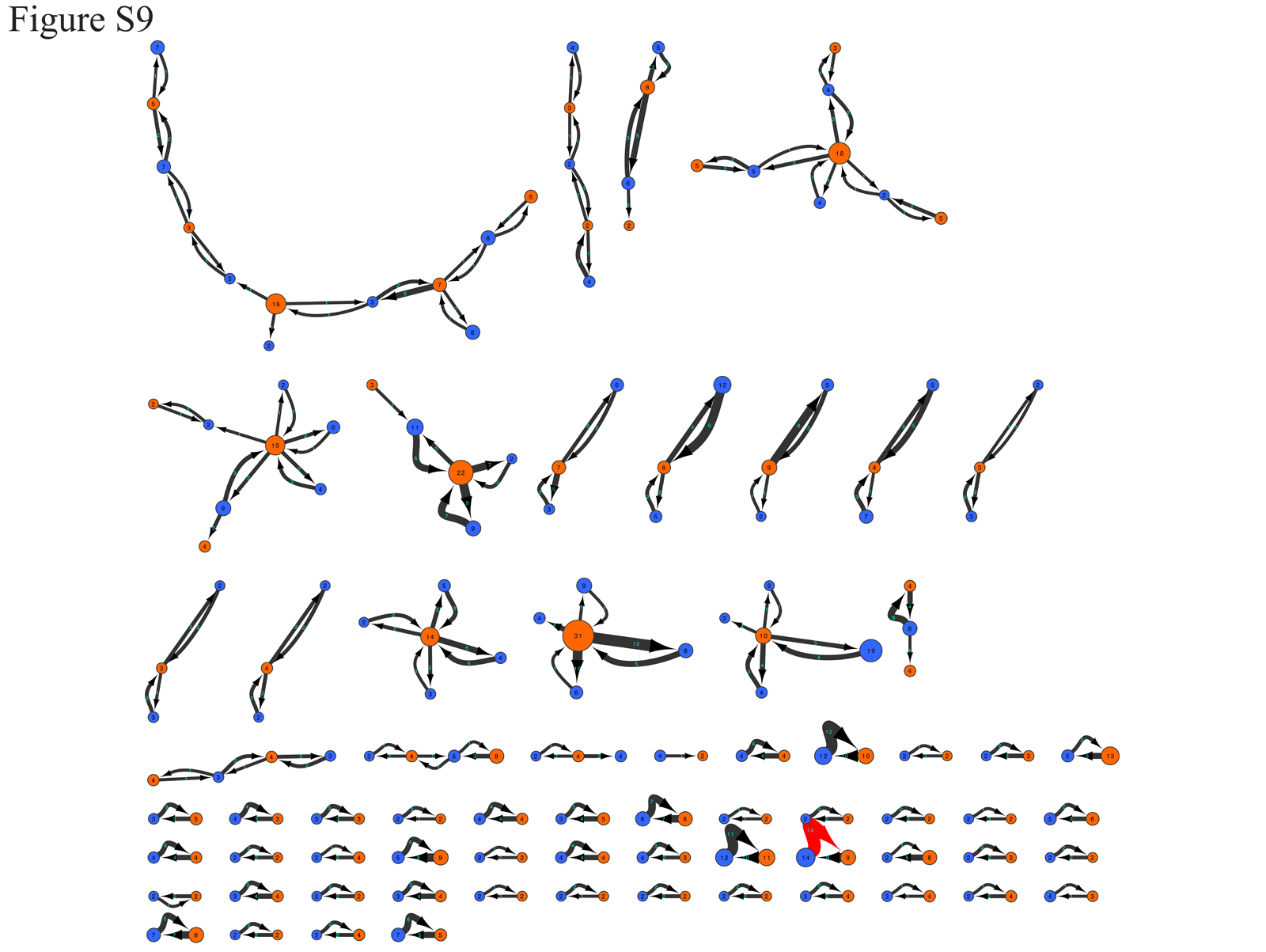
Diagram of conserved cluster between *C. clementina* and *C. sinensis*. The nodes with orange color indicated the clusters of *C. clementina* and the blue ones indicated clusters of *C. sinensis* and size of the nodes were relative to cluster size. The numbers on the edges indicated the number of orthologs could be found from the other species.

**Figure S10.**
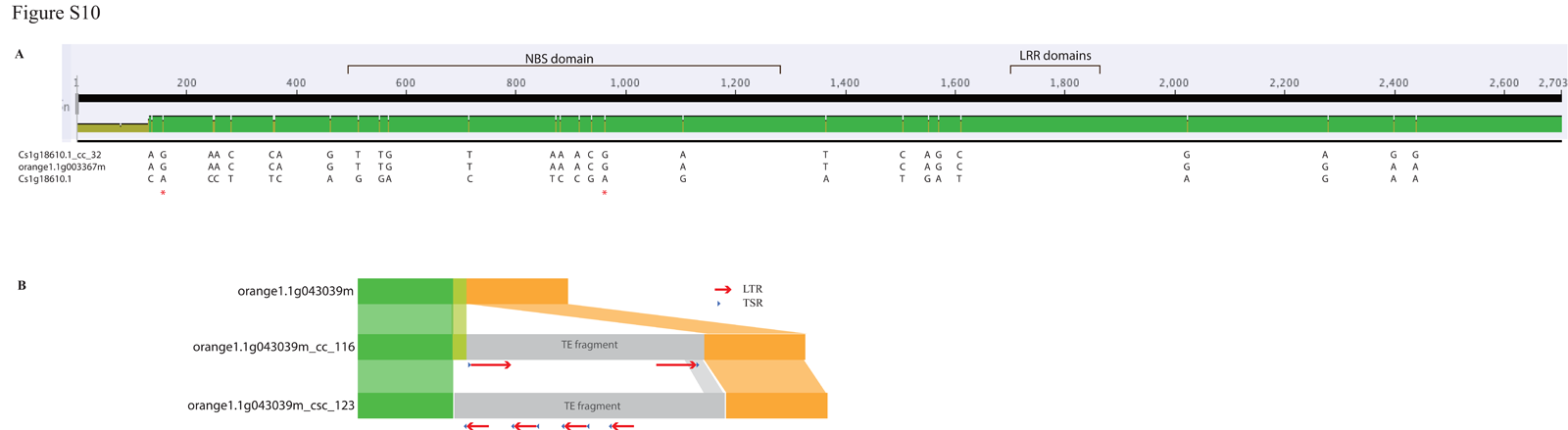
Pseudogenization of NBS-encoding genes of citrus due to mutation (A) and retrotransposon insertion (B). A: Sequences alignment of Cs1g18610.1, Cs1g18610.1_cc_32 and orange1.1g003367m. The bases indicated the variation bases in each citrus species. The star in red color represented stop-codon gaining mutation and may result in pseudogene of Cs1g18610.1. B: LTR retrotransposon insertion in NBS-encoding genes orange1.1g043039m_cc_116 and orange1.1g043039m_csc_123 from Cc and CsCN respectively, and the multiple sequences alignment of the orthologs. LTR, Long terminal repeats; TSR, Target site repeats.

**Figure S11.**
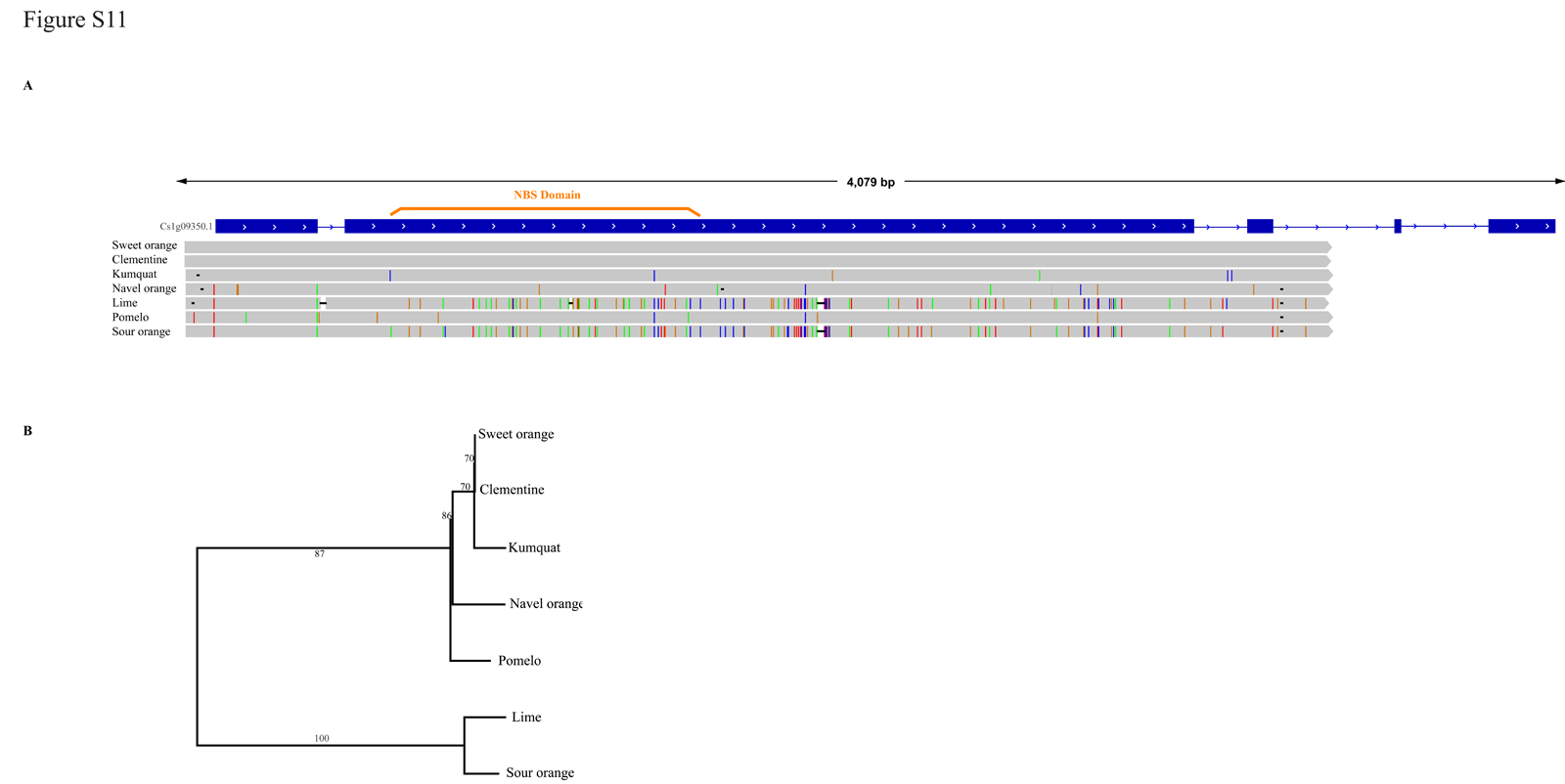
Validation and phylogenetic analysis of NBS-LRR gene, Cs1g09350.1, in different citrus species. A) multiple sequences alignment, the blue bars indicate the exons and the thin lines indicate the introns. The vertical lines in each alignment blocks indicate mutations comparing with the reference sequences from *C. sinensis*. B) neighbor-joining phylogenetic tree of the orthologs of Cs1g09350.1 in different citrus species and the bootstrapping values displayed on the branches.

**Table S1.** The NBS genes in three Citrus genomes

**Table S2.** Classification, cluster and clade of Citrus NBS genes

**Table S3.** LRR domain distribution of citrus NBS gene

**Table S4.** Positive selection sites distribution

**Table S5.** The exome coverage of NBS genes from the 3 clementine and 3 pummelo resequencing genomes

**Table S6.** Citrus NBS gene interapted with LTR transposons

## References

1. Dangl JL, Jones JD: Plant pathogens and integrated defence responses to infection. Nature 2001, 411(6839):826–833.

2. Jia Y, McAdams SA, Bryan GT, Hershey HP, Valent B: Direct interaction of resistance gene and avirulence gene products confers rice blast resistance. The EMBO Journal 2000, 19(15):4004–4014.

3. Ellis JG, Lawrence GJ, Luck JE, Dodds PN: Identification of regions in alleles of the flax rust resistance gene L that determine differences in gene-for-gene specificity. The Plant Cell Online 1999, 11(3):495–506.

4. Ellis J, Dodds P, Pryor T: The generation of plant disease resistance gene specificities. Trends in plant science 2000, 5(9):373–379.

5. Meyers BC, Kozik A, Griego A, Kuang H, Michelmore RW: Genome-wide analysis of NBS-LRR–encoding genes in Arabidopsis. The Plant Cell Online 2003, 15(4):809–834.

6. Kohler A, Rinaldi C, Duplessis S, Baucher M, Geelen D, Duchaussoy F, Meyers BC, Boerjan W, Martin F: Genome-wide identification of NBS resistance genes in Populus trichocarpa. Plant molecular biology 2008, 66(6):619–636.

7. Mun J-H, Yu H-J, Park S, Park B-S: Genome-wide identification of NBS-encoding resistance genes in Brassica rapa. Molecular Genetics and Genomics 2009, 282(6):617–631.

8. Wan H, Yuan W, Bo K, Shen J, Pang X, Chen J: Genome-wide analysis of NBS-encoding disease resistance genes in Cucumis sativus and phylogenetic study of NBS-encoding genes in Cucurbitaceae crops. BMC genomics 2013, 14(1):109.

9. Sun X, Wang G-L: Genome-wide identification, characterization and phylogenetic analysis of the rice LRR-kinases. PloS one 2011, 6(3):e16079.

10. Zhou T, Wang Y, Chen J-Q, Araki H, Jing Z, Jiang K, Shen J, Tian D: Genome-wide identification of NBS genes in japonica rice reveals significant expansion of divergent non-TIR NBS-LRR genes. Molecular Genetics and Genomics 2004, 271(4):402–415.

11. Yang S, Zhang X, Yue J-X, Tian D, Chen J-Q: Recent duplications dominate NBS-encoding gene expansion in two woody species. Molecular Genetics and Genomics 2008, 280(3):187–198.

12. Ameline-Torregrosa C, Wang B-B, O'Bleness MS, Deshpande S, Zhu H, Roe B, Young ND, Cannon SB: Identification and characterization of nucleotide-binding site-leucine-rich repeat genes in the model plant Medicago truncatula. Plant physiology 2008, 146(1):5–21.

13. Pan Q, Wendel J, Fluhr R: Divergent evolution of plant NBS-LRR resistance gene homologues in dicot and cereal genomes. Journal of Molecular Evolution 2000, 50(3):203–213.

14. Michelmore RW, Meyers BC: Clusters of resistance genes in plants evolve by divergent selection and a birth-and-death process. Genome Research 1998, 8(11):1113–1130.

15. Guo Y-L, Fitz J, Schneeberger K, Ossowski S, Cao J, Weigel D: Genome-wide comparison of nucleotide-binding site-leucine-rich repeat-encoding genes in Arabidopsis. Plant physiology 2011, 157(2):757–769.

16. Yang S, Li J, Zhang X, Zhang Q, Huang J, Chen J-Q, Hartl DL, Tian D: Rapidly evolving R genes in diverse grass species confer resistance to rice blast disease. Proceedings of the National Academy of Sciences 2013.

17. Barrett H, Rhodes A: A numerical taxonomic study of affinity relationships in cultivated Citrus and its close relatives. Systematic Botany 1976:105–136.

18. . Scora RW: On the history and origin of Citrus. Bulletin of the Torrey Botanical Club 1975:369–375.

19. Nicolosi E, Deng Z, Gentile A, La Malfa S, Continella G, Tribulato E: Citrus phylogeny and genetic origin of important species as investigated by molecular markers. Theoretical and Applied Genetics 2000, 100(8):1155–1166.

20. Li X, Xie R, Lu Z, Zhou Z: The origin of cultivated citrus as inferred from internal transcribed spacer and chloroplast DNA sequence and amplified fragment length polymorphism fingerprints. Journal of the American Society for Horticultural Science 2010, 135(4):341–350.

21. Xu Q, Chen LL, Ruan X, Chen D, Zhu A, Chen C, Bertrand D, Jiao WB, Hao BH, Lyon MP et al: The draft genome of sweet orange (Citrus sinensis). Nature genetics 2013, 45(1):59–66.

22. Xu Q, Biswas MK, Lan H, Zeng W, Liu C, Xu J, Deng X: Phylogenetic and evolutionary analysis of NBS-encoding genes in Rutaceae fruit crops. Molecular Genetics and Genomics 2011, 285(2):151–161.

23. Meyers BC, Kozik A, Griego A, Kuang H, Michelmore RW: Genome-wide analysis of NBS-LRR-encoding genes in Arabidopsis. The Plant cell 2003, 15(4):809–834.

24. Kohler A, Rinaldi C, Duplessis S, Baucher M, Geelen D, Duchaussoy F, Meyers BC, Boerjan W, Martin F: Genome-wide identification of NBS resistance genes in Populus trichocarpa. Plant molecular biology 2008, 66(6):619–636.

25. Price MN, Dehal PS, Arkin AP: FastTree: computing large minimum evolution trees with profiles instead of a distance matrix. Molecular biology and evolution 2009, 26(7):1641–1650.

26. Plocik A, Layden J, Kesseli R: Comparative analysis of NBS domain sequences of NBS-LRR disease resistance genes from sunflower, lettuce, and chicory. Molecular phylogenetics and evolution 2004, 31(1):153–163.

27. . Li H: Aligning sequence reads, clone sequences and assembly contigs with BWA-MEM. arXiv:13033997v2 2013.

28. Zhou T, Wang Y, Chen JQ, Araki H, Jing Z, Jiang K, Shen J, Tian D: Genome-wide identification of NBS genes in japonica rice reveals significant expansion of divergent non-TIR NBS-LRR genes. Molecular genetics and genomics: MGG 2004, 271(4):402–415.

29. Wang Y, Tang H, Debarry JD, Tan X, Li J, Wang X, Lee TH, Jin H, Marler B, Guo H et al: MCScanX: a toolkit for detection and evolutionary analysis of gene synteny and collinearity. Nucleic acids research 2012, 40(7):e49.

30. Michelmore RW, Meyers BC: Clusters of resistance genes in plants evolve by divergent selection and a birth-and-death process. Genome research 1998, 8(11):1113–1130.

31. Tan S, Wu S: Genome Wide Analysis of Nucleotide-Binding Site Disease Resistance Genes in Brachypodium distachyon. Comparative and functional genomics 2012, 2012:418208.

32. Zhang Z, Carriero N, Zheng D, Karro J, Harrison PM, Gerstein M: PseudoPipe: an automated pseudogene identification pipeline. Bioinformatics 2006, 22(12):1437–1439.

33. Punta M, Coggill PC, Eberhardt RY, Mistry J, Tate J, Boursnell C, Pang N, Forslund K, Ceric G, Clements J et al: The Pfam protein families database. Nucleic Acids Research 2012, 40(D1):D290–D301.

34. Yang S, Feng Z, Zhang X, Jiang K, Jin X, Hang Y, Chen J-Q, Tian D: Genome-wide investigation on the genetic variations of rice disease resistance genes. Plant molecular biology 2006, 62(1-2):181–193.

35. Finn RD, Clements J, Eddy SR: HMMER web server: interactive sequence similarity searching. Nucleic acids research 2011, 39(Web Server issue):W29-37.

36. Punta M, Coggill PC, Eberhardt RY, Mistry J, Tate J, Boursnell C, Pang N, Forslund K, Ceric G, Clements J et al: The Pfam protein families database. Nucleic acids research 2012, 40(Database issue):D290-301.

37. Boeckmann B, Bairoch A, Apweiler R, Blatter MC, Estreicher A, Gasteiger E, Martin MJ, Michoud K, O'Donovan C, Phan I et al: The SWISS-PROT protein knowledgebase and its supplement TrEMBL in 2003. Nucleic acids research 2003, 31(1):365–370.

38. Altschul SF, Madden TL, Schaffer AA, Zhang J, Zhang Z, Miller W, Lipman DJ: Gapped BLAST and PSI-BLAST: a new generation of protein database search programs. Nucleic acids research 1997, 25(17):3389–3402.

39. Birney E, Clamp M, Durbin R: GeneWise and Genomewise. Genome research 2004, 14(5):988–995.

40. Moreno-Hagelsieb G, Latimer K: Choosing BLAST options for better detection of orthologs as reciprocal best hits. Bioinformatics 2008, 24(3):319–324.

41. Suyama M, Torrents D, Bork P: PAL2NAL: robust conversion of protein sequence alignments into the corresponding codon alignments. Nucleic acids research 2006, 34(Web Server issue):W609-612.

42. Yang Z: PAML 4: phylogenetic analysis by maximum likelihood. Molecular biology and evolution 2007, 24(8):1586–1591.

43. Katoh K, Standley DM: MAFFT Multiple Sequence Alignment Software Version 7: Improvements in Performance and Usability. Molecular biology and evolution 2013, 30(4):772–780.

44. Prosperi MC, Ciccozzi M, Fanti I, Saladini F, Pecorari M, Borghi V, Di Giambenedetto S, Bruzzone B, Capetti A, Vivarelli A et al: A novel methodology for large-scale phylogeny partition. Nature communications 2011, 2:321.

45. Delorenzi M, Speed T: An HMM model for coiled-coil domains and a comparison with PSSM-based predictions. Bioinformatics 2002, 18(4):617–625.

46. Lupas A: Prediction and analysis of coiled-coil structures. Methods in enzymology 1996, 266:513–525.

47. Bailey TL, Boden M, Buske FA, Frith M, Grant CE, Clementi L, Ren J, Li WW, Noble WS: MEME SUITE: tools for motif discovery and searching. Nucleic acids research 2009, 37(Web Server issue):W202-208.

48. Xu Z, Wang H: LTR_FINDER: an efficient tool for the prediction of full-length LTR retrotransposons. Nucleic acids research 2007, 35(Web Server issue):W265-268.

49. Sawyer S: Statistical tests for detecting gene conversion. Molecular biology and evolution 1989, 6(5):526–538.

50. Quinlan AR, Hall IM: BEDTools: a flexible suite of utilities for comparing genomic features. Bioinformatics 2010, 26(6):841–842.

51. Zhou L, Powell CA, Hoffman MT, Li W, Fan G, Liu B, Lin H, Duan Y: Diversity and plasticity of the intracellular plant pathogen and insect symbiont “Candidatus Liberibacter asiaticus” as revealed by hypervariable prophage genes with intragenic tandem repeats. Applied and environmental microbiology 2011, 77(18):6663–6673.

